# Protein aggregation and calcium dysregulation are the earliest hallmarks of synucleinopathy in human midbrain dopaminergic neurons

**DOI:** 10.1101/2022.10.28.514238

**Authors:** Gurvir S Virdi, Minee L Choi, James R Evans, Zhi Yao, Dilan Athauda, Stephanie Strohbuecker, Anna I Wernick, Haya Alrashidi, Daniela Melandri, Jimena Perez-Lloret, Plamena R Angelova, Sergiy Sylantyev, Simon Eaton, Simon Heales, Tilo Kunath, Mathew H Horrocks, Andrey Y Abramov, Rickie Patani, Sonia Gandhi

## Abstract

Mutations in the *SNCA* gene cause autosomal dominant Parkinson’s disease (PD), with progressive loss of dopaminergic neurons in the substantia nigra, and accumulation of aggregates of α-synuclein. However, the sequence of molecular events that proceed from the *SNCA* mutation during development, to its end stage pathology is unknown. Utilising human induced pluripotent stem cells (hiPSCs) with *SNCA* mutations, we resolved the temporal sequence of pathophysiological events that occur during neuronal differentiation in order to discover the early, and likely causative, events in synucleinopathies. We adapted a small molecule-based protocol that generates highly enriched midbrain dopaminergic (mDA) neurons (>80%). We characterised their molecular identity using single-cell RNA sequencing and their functional identity through the synthesis and secretion of dopamine, the ability to generate action potentials, and form functional synapses and networks. RNA velocity analyses confirmed the developmental transcriptomic trajectory of midbrain neural precursors into mDA neurons using our approach, and identified key driver genes in mDA neuronal development. To characterise the synucleinopathy, we adopted super-resolution methods to determine the number, size and structure of aggregates in *SNCA*-mutant mDA neurons. At one week of differentiation, prior to maturation to mDA neurons of molecular and functional identity, we demonstrate the formation of small aggregates; specifically, β-sheet rich oligomeric aggregates, in *SNCA*-mutant midbrain immature neurons. The aggregation progresses over time to accumulate phosphorylated aggregates, and later fibrillar aggregates. When the midbrain neurons were functional, we observed evidence of impaired physiological calcium signalling, with raised basal calcium, and impairments in cytosolic and mitochondrial calcium efflux. Once midbrain identity fully developed, *SNCA*-mutant neurons exhibited bioenergetic impairments, mitochondrial dysfunction and oxidative stress. During the maturation of mDA neurons, upregulation of mitophagy and autophagy occured, and ultimately these multiple cellular stresses lead to an increase in cell death by six weeks post-differentiation. Our differentiation paradigm generates an efficient model for studying disease mechanisms in PD, and highlights that protein misfolding to generate intraneuronal oligomers is one of the earliest critical events driving disease in human neurons, rather than a late-stage hallmark of the disease.

## Introduction

Parkinson’s disease (PD) is a progressive neurodegenerative disease primarily characterised by the loss of dopaminergic neurons in the substantia nigra of the midbrain ^1^. Studying the mechanisms that contribute to disease is of immense importance, but has been somewhat limited by a paucity of relevant human models. The development of human-induced pluripotent stem cells (hiPSCs) from patients with familial forms of PD has vastly improved our capability to model and study PD. Taking this advance further, directed differentiation strategies to generate the relevant cell type from the relevant brain region, should allow us to discover mechanisms of disease and define the basis for selective cellular vulnerability. For PD, amongst the earliest and most vulnerable cell populations is the midbrain dopaminergic neuron, and therefore modelling disease with precision requires the ability to robustly generate this neuronal type.

Several protocols have transformed our ability to generate midbrain dopaminergic (mDA) neurons ^2–5^: these mimic *in vivo* developmental programmes, inducing neural induction via dual-SMAD inhibition ^6^, activating sonic hedgehog (Shh), Wnt, and fibroblast growth factor 8 (FGF8) signalling through the use of small-molecules and/or recombinant morphogens. The resulting cells are molecularly defined as midbrain-specific, based on the expression of key genes and proteins, as well as functional and physiological characteristics resembling *in vivo* mDA neurons ^7^. The efficiency of mDA neuron production varies between current differentiation protocols ^8^, and cellular heterogeneity can be challenging when modelling disease. Against this background, we adapted well-established protocols to obtain highly enriched mDA neurons, and we tested their capacity to reliably model key aspects of PD pathogenesis as well as identify early phenotypes.

The pathological hallmark of PD is the presence of insoluble aggregated forms of the protein α - synuclein, encoded by the *SNCA* gene ^9^. Modelling synucleinopathy *in vitro* may therefore be achieved by utilising hiPSC lines with *SNCA* mutations. A range of mutations and multiplications have been discovered in α-synuclein, including the p.A53T point mutation and the triplication of the *SNCA* locus. Both of these genetic changes cause early onset and rapidly progressive form of PD ^10–12^. HiPSC-derived models harbouring *SNCA* mutations have highlighted a range of cellular phenotypes that may lead to cell vulnerability, including mitochondrial dysfunction ^13–15^, ER stress ^16^, lysosomal dysfunction ^17^, oxidative stress, and calcium dysregulation ^18,19^. Nonetheless, disease models are dominated by multiple forms of cellular stress, and it has not been possible to determine the emergence of cellular dysfunction, and resolve the events spatially and temporally.

In this study, we established a modified approach to obtain highly enriched mDA neurons and we demonstrate their capacity to faithfully model key aspects of PD pathogenesis. Applying this method with disease-based modelling we were able to temporally sequence the pathological events in *SNCA*-mutant lines, and delineate the critical events in the midbrain dopaminergic neuron.

## Results

### Highly enriched midbrain dopaminergic neurogenesis from iPSCs

To generate mDA neurons from iPSCs we induced neural conversion using dual-SMAD inhibition ^6^ followed by lineage restriction towards the midbrain using small molecule agonists of Shh and Wnt, purmorphamine and CHIR99021 respectively (Figure 1A). After 14 days, we confirmed mDA neural precursor cell (NPC) identity using RT-PCR for LMX1A, FOXA2, and EN1 (p < 0.005) (Figure 1B) and immunocytochemistry (ICC) for LMX1A, FOXA2 and OTX2 (Figure 1C). Crucially, high co-expression of all three markers, LMX1A, FOXA2 and OTX2, confirmed an enriched culture of mDA NPCs (Ctrl 1: 83% ± 3.5, Ctrl 2: 81.4% ± 2.4, Ctrl 3: 88.7% ± 3.1, Ctrl 4: 88.4% ± 2.2) (Figure 1D). After midbrain patterning, NPCs were maintained in culture for 4 days before terminal differentiation to increase their confluency. Neuronal differentiation was induced using a Notch pathway inhibitor and a Rho-associated protein kinase (ROCK) inhibitor, encouraging cell cycle exit and cell survival respectively. Maturation of mDA neurons was assessed during terminal differentiation and demonstrated an increase of TH-positive cells (Figure 1E). We showed that over 75% of TH-positive cells were obtained after 3 weeks of differentiation using ICC (Ctrl 1: 82.1% ± 3.8, Ctrl 2: 73.1% ± 6.9, Ctrl 3: 77.9% ± 3.2) (Figure 1F & G). Flow cytometry also confirmed that the cultured cells expressed approximately 88% expression of both TH and β-III Tubulin (88.8% ± 1.9, Figure 1H & I, Supplementary Figure 1A & B). Overall, the data confirm high efficiency of neuronal differentiation, specifically into TH-positive neurons in 80-90% of the whole culture, with comparable efficiencies across three control hiPSC lines. To verify the maturity of the TH-positive mDA neurons, we performed qPCR for specific genes including *NR4A2* (Nurr1ļ *KCNJ6* (GIRK2), and *SLC6A3* (DAT) at 3-4-week-old terminally differentiated mDA neurons. qPCR results showed an increase in the expression of these transcriptional markers of terminally differentiated mDA neurons relative to midbrain NPCs (Supplementary Figure 1C). ICC quantification using a validated GIRK2 antibody also confirmed the expression of GIRK2 in mDA neurons which further supports the midbrain identity (Supplementary Figure 1D), given that GIRK2 is highly expressed in dopaminergic neurons in the A9 region of the midbrain (typical of the substantia nigra).

**Figure 1.**
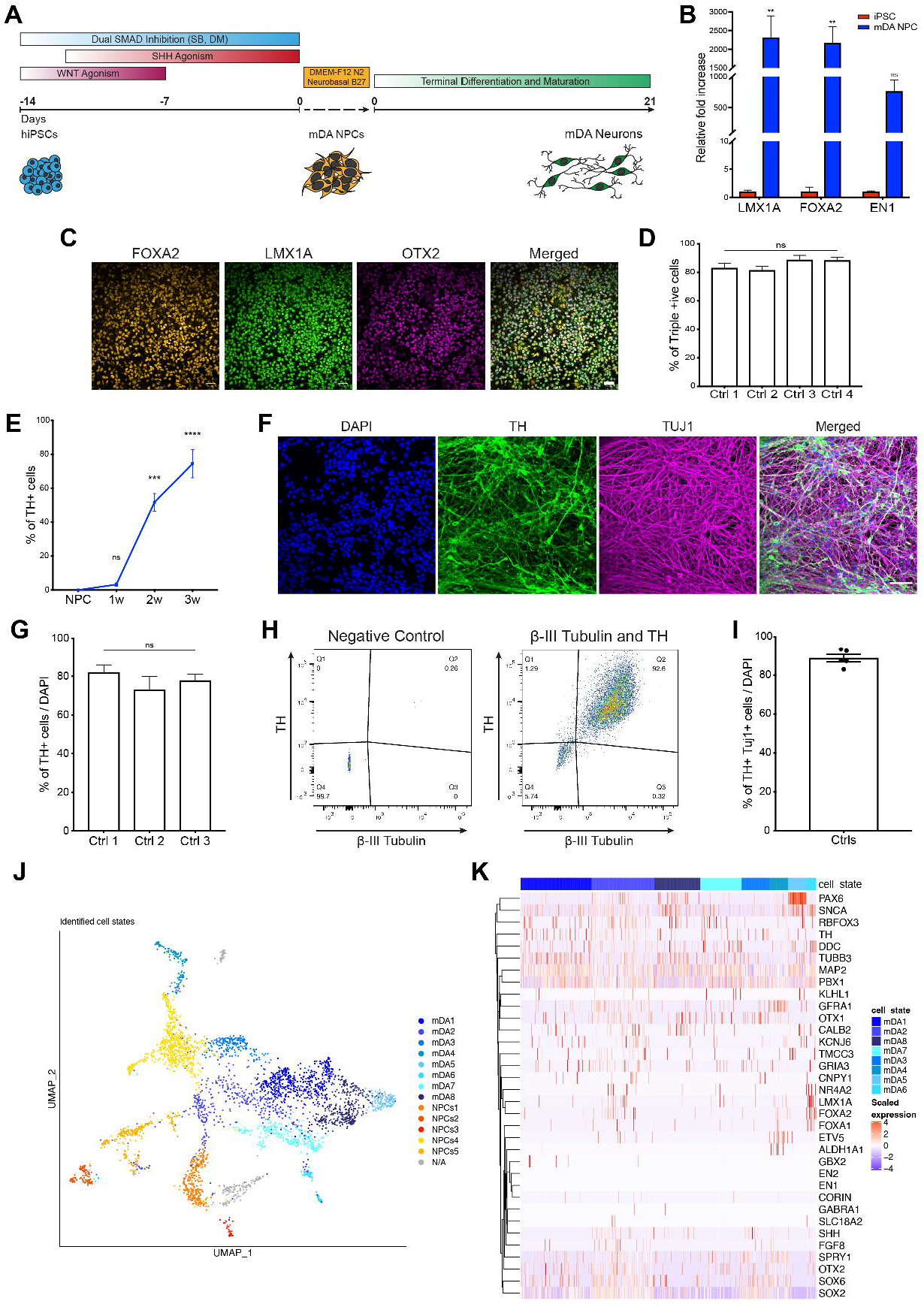
Characterisation of an enriched population of mDA NPCs and neurons. **(A**) Differentiation protocol to generate mDA neurons. **(B)** Quantitative PCR showing an increased mRNA for mDA NPC markers LMX1A, FOXA2, and EN1 relative to hiPSCs (n= 3 different lines across 4 independent neuronal inductions, ns p > 0.05 ** p < 0.005, ordinary twoway ANOVA). Data plotted as ±SEM. **(C)** Representative ICC images showing high expression of mDA NPC markers FOXA2, LMX1A, and the forebrain/midbrain marker OTX2 (scale bar = 50μm). **(D)** Quantification of immunocytochemistry images showing >80% of cells co-express mDA NPC markers FOXA2, LMX1A, and OTX2 (n = 4 different lines across 3 independent neuronal inductions, ns p > 0.05, ordinary one-way ANOVA). Values plotted as ±SEM. **(E)** Quantification of ICC images showing the increase in expression of the mDA mature neuron marker, TH, as the neurons differentiate and mature (n = 3 different lines across 1 independent neuronal induction, ns p > 0.05, *** p < 0.0005, **** p < 0.0001, ordinary one-way ANOVA). Values plotted as ±SEM. **(F)** Representative ICC images showing TH and TUJ1 expression after 3 weeks of differentiation (41 days from hiPSC) (scale bar = 50μm). **(G)** Quantification of ICC images after 3 weeks of differentiation showing approximately 80% cells express TH (n= 3 different lines across 3 independent neuronal inductions, ns p > 0.05, ordinary one-way ANOVA). Values plotted as ±SEM. **(H)** Representative dot plots of single-cell suspensions showing % TH and β-III Tubulin +ve cells 3 weeks after terminal differentiation. A negative control (DAPI only) was used to determine quantification thresholds to set gating (n = 10,000 events recorded per measurement). **(I)** Quantification of flow cytometry analysis showing over 80% of DAPI positive cells co-express TH and β-III Tubulin (n = 3 lines across 2 independent neuronal inductions). Values plotted as ±SEM. **(J)** A UMAP plot showing the 14 clusters identified from single-cell RNA-seq after 4 weeks of differentiation. Neuronal mDA (mDA1-8) clusters are in blue, and NPC (NPCs1-5) is in red, orange and yellow. An unidentified cluster (N/A) is coloured in grey. **(K)** Heat map showing expression of genes in clusters identified as mDA neurons (mDA1-8) in the culture after 4 weeks of differentiation. Each line represents a cell from that cluster. Genes annotated in red correspond to mDA NPC markers, and those in black correspond to mDA neuron markers.

We performed single-cell RNA-seq on 4 week terminally differentiated mDA neurons. The cultured cells were clustered into identities based on their transcriptional profiles. Clustering of all grouped control mDA neurons revealed 13 clusters, of which seven expressed key mDA neuron genes, confirming their identity as mDA neurons (Figure 1J). The key mDA genes expressed include GDNF receptor (*GFRA1*), *SOX6, PBX1, TH, KCNJ6, NR4A2*, and *DDC*, suggesting that the majority of captured cells were of mDA identity, and were abundant within the culture (Figure 1K) (Supplementary Figure 1G). We also detected several clusters which were similar to mDA NPCs, or early neurons (5 clusters) (Figure 1J). These clusters expressed key NPC genes e.g *FOXA2, LMX1A, OTX2, FGF8, and SHH*, as well as mDA neuronal markers including *GFRA1, DDC, NR4A2, TH*, and *KCNJ6* (Supplementary Figure 1H & I, Supplementary Figure 2). In addition, most clusters expressed *SNCA*, highlighting the potential to use the protocol to model synucleinopathy (Supplementary Figure 1F).

In order to characterise the lineage relationship between different cell clusters during cellular differentiation, we performed RNA-velocity analysis, scVelo ^20^ in single-cell RNA-seq data on 4-week-old mDA neuronal cultures ^21^. RNA velocity, i.e., the change in RNA abundance for each gene in a single cell, can be used to predict the future transcriptomic state of a cell, and thus infer developmental trajectories. We confirmed the expected neuronal differentiation trajectory from the NPC clusters towards mature mDA neurons (Figure 2A & B). In addition, we found that several mDA neuron clusters were on a different differentiation trajectory to other mDA neuron clusters (Figure 2C). The dynamical model used by scVelo allowed us to recover latent time, which approximates the cell’s internal clock. Using latent time to reconstruct the temporal sequence of cellular fates, we observed that the NPC cell clusters were earlier than mDA neurons and were spread across latent time, suggesting different clusters were at different points on the path to form mature mDA neurons (Figure 2Di). We display the dynamic behaviour of putative driver genes for each cluster that were systematically detected by likelihoods in the dynamical model, where the highest scored genes associated with neuronal differentiation include *STMN2, DCX*, and *SYT1*, and the highest scored genes for NPC clusters include *OTX2, LMX1A*, and *QKI* (Figure 2Dii, Supplementary Figure 3B). In addition, each NPC cluster had a different confidence probability showing what mDA neuronal cluster trajectory path they were predicted to be on to (Supplementary Figure 3C). Lastly, we performed gene ontology (GO) enrichment ^22^ for the top 100 driver genes for each cluster. The driver genes in NPC clusters were associated with GO terms related to proliferation, as well as neuronal differentiation (Figure 2Ei & ii). The mDA neuron clusters, on the other hand, showed enrichment in GO terms associated with complex synapse formation, as well as ion transport development (Figure 2Fi & ii).

**Figure 2.**
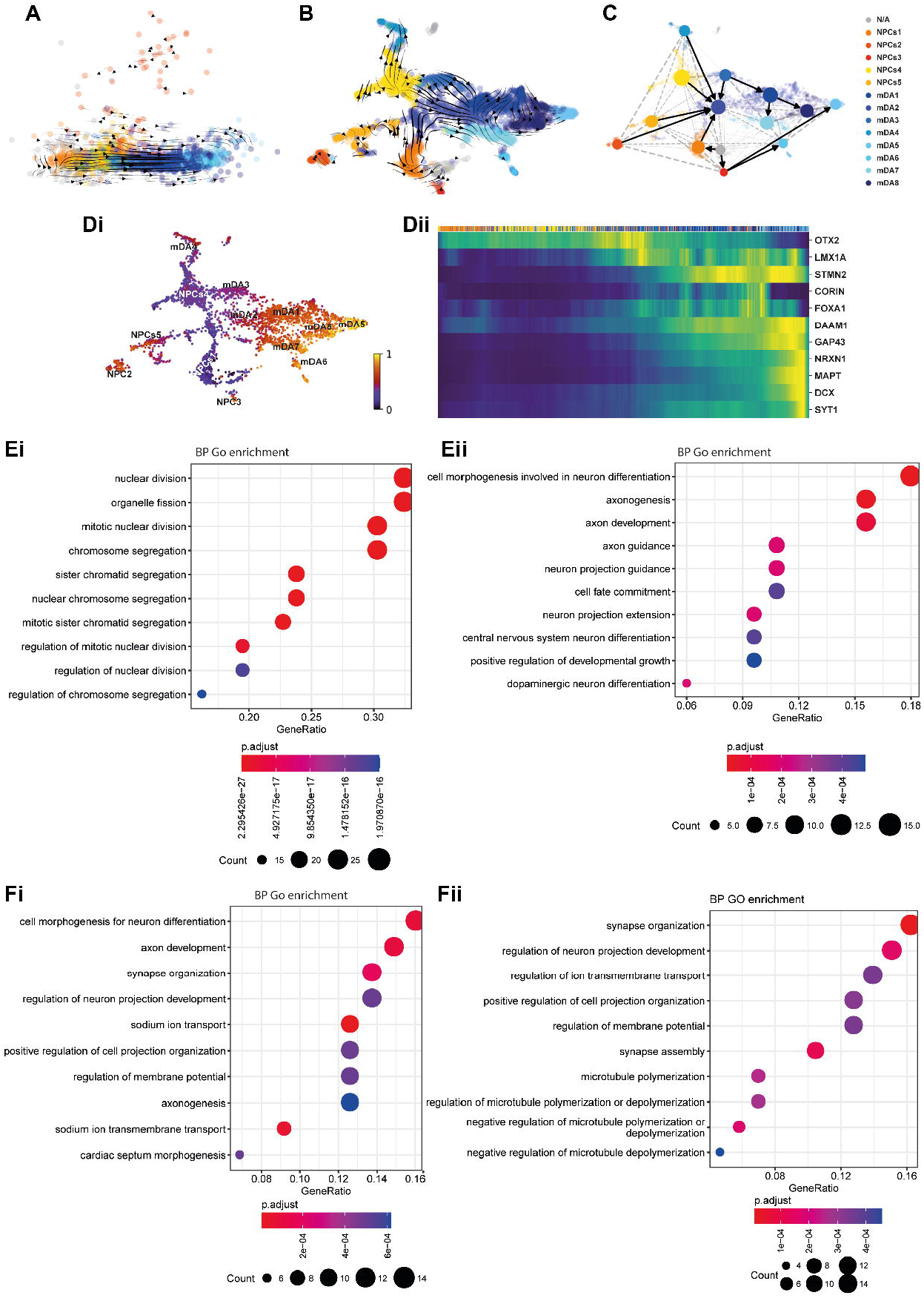
RNA-velocity demonstrates developmental streams from NPCs into mDA neurons reminiscent of a developing midbrain. **(A)** PCA visualises the progression from more precursor cells (NPCs) (yellow-red colours) towards more mDA neuronal cells (blue colours). RNA-velocity trajectories are indicated by arrows annotated with cell type clusters. **(B)** Velocity stream visualised in UMAP reveals a more detailed progression of the direction of differentiation for each cluster (NPCs in yellow-red, and mDA clusters in blue). **(C)** PAGA graph showing the NPC cluster transition into mDA neuron clusters. **(Di & ii)** Velocity inferred latent time analysis. **(Di)** Latent time visualised in UMAP, showing mDA clusters are more later (brighter colour) in time than NPC clusters (darker colours). **(Dii)** A heatmap listing a selection of cluster driver genes sorted according to their inferred latent time. NPC genes arise earlier in time, and mDA neuron genes arise later in latent time. **(E & F)** GO enrichment analysis to clarify gene categories per cell cluster based on the functional characteristics of the driver genes. **(Ei & ii)** Example of GO enrichment of two NPC clusters showing that **(Ei)** some clusters are more proliferative, and some **(Eii)** are more differentiated and on the neuronal pathway. **(Fi & ii)** Example of GO enrichment of two mDA neuron clusters showing complex neuronal pathways which are highly activated.

Our analysis demonstrates that, at 28 days post-differentiation, our method predominantly generates cells of mDA neuronal identity, with a clear developmental trajectory defined for the remaining NPCs to differentiate into mDA neurons.

### hiPSC-derived mDA neurons exhibit functional neuronal and dopaminergic specialisation

In order to test the functional properties of the mDA neurons, we first determined whether they expressed voltage-dependent calcium channels (VDCCs) that are expressed primarily in neurons. Opening of calcium channels was induced using a high concentration of KCl (50mM), which depolarises the plasma membrane, opening VDCCs, resulting in a large influx of calcium into the neurons. A cytosolic Ca^2+^ signal was induced by KCl in all cultures, in approximately 65% of cells after 2 weeks, increasing to over 85% after 5 weeks of differentiation (2 weeks = 65.5% ± 5.7, 3 weeks = 73.1% ± 2.3, 4 weeks = 73.2 ± 1.7, 5 weeks = 88.6% ± 1.8, p = 0.0075) (Figure 3A & B, Supplementary Figure 4A). After 3 weeks of differentiation, we observed spontaneous calcium fluctuations in neurons (Figure 3C), which are a key hallmark of mDA dopaminergic neurons in the substantia nigra and contributes to their pace-making characteristic ^23^.

**Figure 3.**
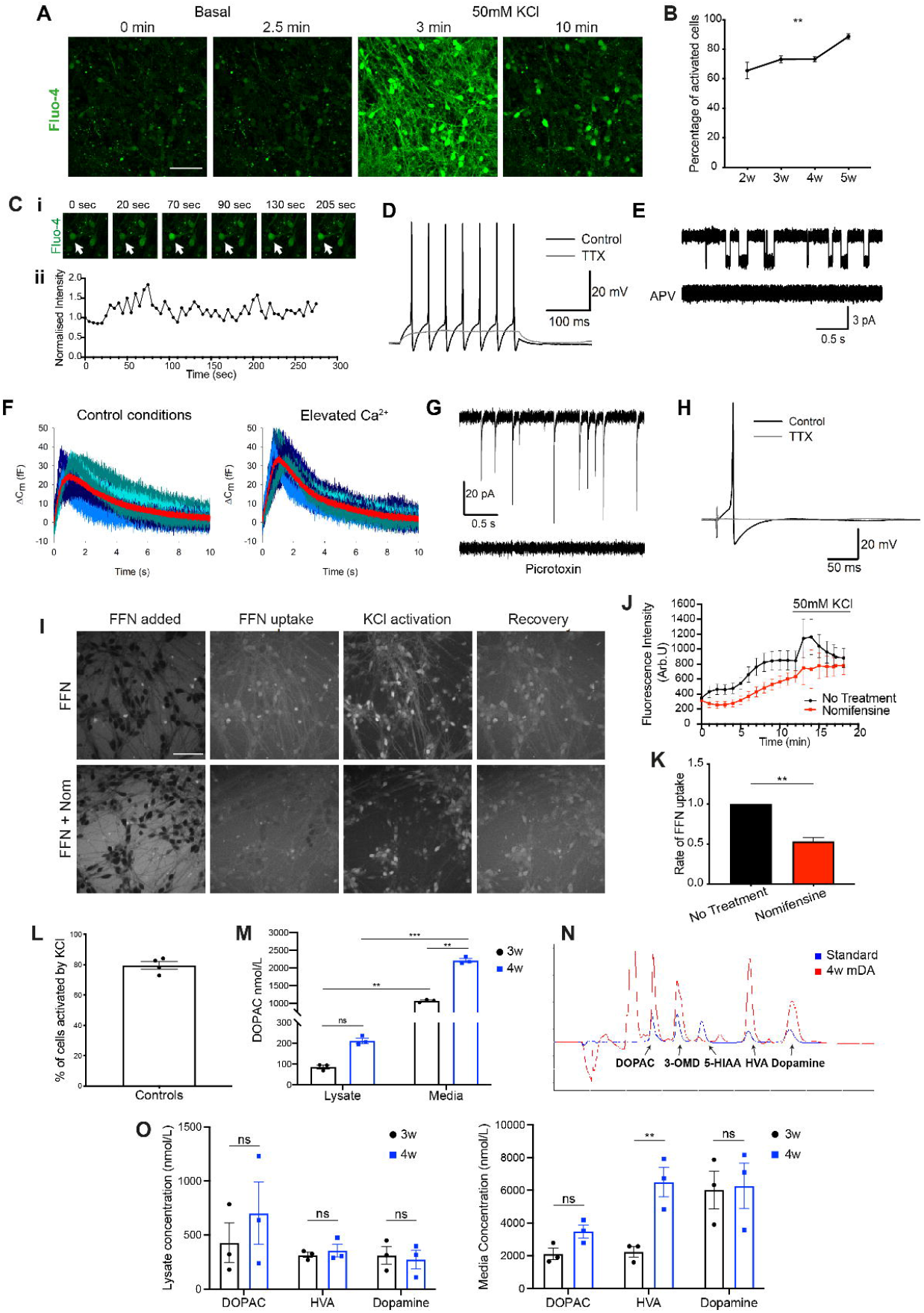
Functional characterisation of mDA neurons. **(A)** Representative time series images of mDA neurons at 5 weeks of terminal differentiation, in response to KCl (scale bar = 50μm). **(B)** Quantification of number of cells with calcium response (n = 3 separate lines across 1 independent neuronal induction, ns p > 0.05, ** p = 0.0075, twoway ANOVA). Values plotted as ±SEM. **(C) (Ci)** Representative time series images of spontaneous calcium activity at 3 weeks of terminal differentiation, arrow indicates highlighted cell **(Cii)** trace of Fluo-4 intensity in highlighted cell. **(D)** APs triggered by step current injection in mDA neurons at day 30 of differentiation. 1 mM tetrodotoxin (TTX) fully suppresses APs, confirming involvement of voltage-gated sodium channels. **(E)** Single-channel openings of NMDA receptors in an outside-out patch excised from mDA neurons at day 30 of differentiation. Top trace: application of 10 mM glutamate + 10 mM glycine triggers single-channel openings with two different conductance states. Bottom trace: 50 mM APV fully suppresses effect of glutamate + glycine, confirming pharmacological profile of NMDA receptors. **(F)** Changes in whole-cell membrane capacitance evoked by stimulation series confirm elevated intensity of vesicle release in mDA neurons at day 70 of differentiation. Shadows of blue, high noise: 10 consequent individual traces; superficial red trace with low noise: averaged trace. Left: control. Right: elevated Ca^2+^ magnifies the effect of external stimulation on membrane capacitance, thus confirming involvement of presynaptic Ca^2+^-dependent mechanism of vesicle release. **(G)** Spontaneous postsynaptic activity confirms presence of functional synapses in mDA neurons at day 70 of differentiation. Top trace: spontaneous postsynaptic currents. Bottom trace: 50 mM of picrotoxin fully suppresses spontaneous synaptic activity, confirming its generation by GABA_A_ receptors. **(H)** AP generation in response to field stimulation confirms establishment of a functional neuronal network in mDA neurons at day 105 of differentiation. **(I)** Representative images showing uptake of the DAT fluorescent substrate FFN102 (FFN) in mDA neurons. First panel shows FFN dye outside the cells. Second panel shows FFN inside cells. Third panel shows the response of FFN upon stimulation with 50mM KCl and the recovery in the fourth panel. Lower panels show FFN uptake in the presence of the DAT inhibitor, nomifensine (scale bar = 50μm). **(J)** Representative trace showing increase in intracellular FFN with time, as well as response to KCl stimulation. Nomifensine reduces the rate of FFN uptake (n = 15-20 cells per condition). Values plotted as ±SD. **(K)** Quantification of the normalised rate of FFN uptake (n = 3 separate lines over 3 independent neuronal inductions, Welch’s t-test, ** p = 0.0024). Values plotted as ±SEM. **(L)** Quantification of number of FFN positive cells once stimulated by KCl (n = 3 separate lines, 3 independent neuronal inductions). Values plotted as ±SEM. **(M)** Quantification of the amount (nmol/L) of the metabolite DOPAC in basal 3- and 4-week-old mDA neurons (n = 3 separate lines across 1 neuronal induction, ns p > 0.05, ** p < 0.008, *** p = 0.0002, ordinary two-way ANOVA). Values plotted as ±SEM. **(N)** Example chromatogram from a 4-week-old mDA neuron culture treated with L-Dopa (red trace) against a standard (blue trace) showing the presence or absence (peaks) of the metabolites DOPAC, 3-O-Methyldopa (3-OMD), 5-HIAA, HVA, Dopamine. **(O)** Quantification of the (nmol/L) metabolites DOPAC, HVA, and Dopamine in lysate, or media of L-Dopa treated 3- and 4-week-old mDA neurons (n = 3 separate lines across 1 neuronal induction, ns p > 0.05, ** p < 0.005, ordinary two-way ANOVA). Values plotted as ±SEM.

We then investigated the functional electrophysiological characteristics of the mDA neurons by examining single-channel, cells and network effects of the culture. We confirmed the generation of action potentials (AP) activated by a depolarizing voltage step, involving the classical tetrodotoxin (TTX)-sensitive voltage-gated sodium channel (Figure 3D). We confirmed the presence of excitatory NMDA receptors, shown by the triggered single-channel openings in outside-patches pulled from cell soma upon application of 10 mM glutamate + 10 mM glycine, and fully inhibited by further application of 50 mM APV, a selective antagonist of NMDA receptors (Figure 3E). Then, to examine whether the mDA neurons had a functional presynaptic neurotransmitter release system, we recorded changes in cell membrane capacitance (Cm) in response to a field stimulation train of five stimuli with 50 ms intervals. We tested the involvement of the classical Ca^2+^-dependent mechanism of presynaptic vesicle release by repeating the experiment under normal (2 mM) and elevated (5 mM) Ca^2+^ concentration. We observed a clear increase between C_m_ value under normal Ca^2+^ concentration and after an elevated Ca^2+^ concentration (Figure 3F), confirming Ca^2+^-dependent presynaptic vesicle release. Next we performed a whole-cell voltage-clamp recording of spontaneous postsynaptic activity. We observed postsynaptic currents, which were suppressed by 50 mM of the selective antagonist of GABAA receptor, picrotoxin (Figure 3G). This confirmed the presence of the main type of inhibitory neuroreceptors (GABAA receptors) and a functional postsynaptic system with a classical response to spontaneously released GABA. Lastly, we tested whether our mDA neurons could establish a functional neural network by performing whole-cell current-clamp recordings, delivering field electrical stimuli in a perfusion chamber. These stimuli generated APs of classical shape, confirming the presence of interneuronal crosstalk (Figure 3H). Thus, we show that the mDA neurons display many electrophysiological characteristics that confirm their behaviour as neurons forming functional networks.

The function of the dopamine transporter (DAT) can be assessed by measuring the uptake of the fluorogenic DAT substrate FFN102 (FFN) ^24^. When incubated with FFN, followed by a wash, the dye can be seen inside mDA neurons (Supplementary Figure 4B). The uptake of FFN in neurons is further activated upon KCl induced calcium influx in the majority of neurons (Figure 3I & L) (cells activated by KCl = 79.5% ± 2.5). Inhibition of the DAT using the compound Nomifensine resulted in a significantly reduced uptake rate of FFN into neurons (No treatment = 1.00 ± 0.00, Nomifensine = 0.53 ± 0.05, p = 0.0024) (Figure 3K) (Supplementary Figure 4C), as well as a reduced KCl stimulated uptake (Figure 3J). This data suggests that the mDA neurons exhibit functional dopamine transport mechanisms.

Lastly, we used HPLC with electrochemical detection to investigate dopamine metabolism in the mDA neurons. Under basal conditions, mDA neurons released dopamine metabolite 3,4-Dihydroxyphenylacetic acid (DOPAC), which increased with time *in vitro* (Figure 3M) (Lysate: 3w = 86.5 nmol/L ± 9, 4w = 212.6 nmol/L ± 13 p > 0.05; Media: 3w = 1072 nmol/L ± 24, 4w = 2212 nmol/L ± 56, p = 0.0041). Therefore, the cells were able to synthesise and metabolise dopamine under basal conditions. Following pre-treatment of the mDA neurons with the dopamine precursor, L-3,4-dihydroxyphenylalanine (L-Dopa), we detected the presence of dopamine, and its metabolites (DOPAC, homovanillic acid [HVA]) in 3- and 4-week terminally differentiated neurons (Figure 3M). The lack of detection of the main serotonin metabolite 5-hydroxyindoleacetic acid (5-HIAA), usually present in neuronal cells that show mixed dopaminergic/serotonergic characteristics ^25^, suggests the mDA neurons are specifically dopaminergic in nature (Figure 3N). Quantification showed that all metabolites were higher in the media compared to cell lysate (Figure 3O), with the metabolites DOPAC and HVA being present in the highest amounts in the media of 4-week-old mDA neurons (DOPAC = 3493 nmol/L ± 401, HVA = 6494 nmol/L ± 898), and the levels of dopamine being consistent between 3- and 4-week-old neurons (Lysate: 3w = 312 nmol/L ± 80, 4w = 274 nmol/L ± 12.8; Media: 3w = 6029 nmol/L ± 1153, 4w = 6277 nmol/L ± 1387, p > 0.05). Therefore, mDA neurons as early as 3 weeks of differentiation had a functional dopamine metabolism and secretion.

### A human PD model can be established by differentiating patient-derived hiPSCs with *SNCA* mutations into functional mDA neurons

Mutations in *SNCA* cause an autosomal dominant form of PD (*SNCA*-PD). HiPSCs from patients with a p.A53T mutation, triplication of the *SNCA* locus (*SNCA* x3), their isogenic pairs and healthy donors were differentiated into mDA neurons using the protocol described above. All three mutant lines were successfully differentiated into mDA neurons, highlighted by the high enrichment of TH and MAP2 protein after 3 weeks of differentiation (Ctrl = 87.9% ± 2.7 TH-positive, 96.2% ±1.6 MAP2-positive, A53T = 83.1% ± 5.3 TH-positive, 89.3% ± 4.7 MAP2-positive, SNCA x3 = 76.2% ± 5.8 TH-positive, 96.0% ± 1.2 MAP2-positive) (Figure 4A & B). Using RT-qPCR, we first found a 3-4-fold increase in *SNCA* mRNA in the *SNCA* x3 line compared to the control lines at 3 weeks of differentiation (Figure 4C). The A53T lines exhibited a similar amount of *SNCA* mRNA compared to the control lines (Ctrl = 1.0 ± 0.13, A53T = 1.4 ± 0.04, SNCA x3 = 3.6 ± 0.5, p < 0.0001). Using an antibody that recognises total α-synuclein, we detected α-synuclein protein expression across all lines at 4 weeks of differentiation, with a significant increase in the expression of α-synuclein in the *SNCA* x3 line (Figure 4D & E) (Ctrl = 1.00 ± 0.03, A53T = 1.05 ± 0.02, SNCA x3 = 1.24 ± 0.06, p < 0.005).

**Figure 4.**
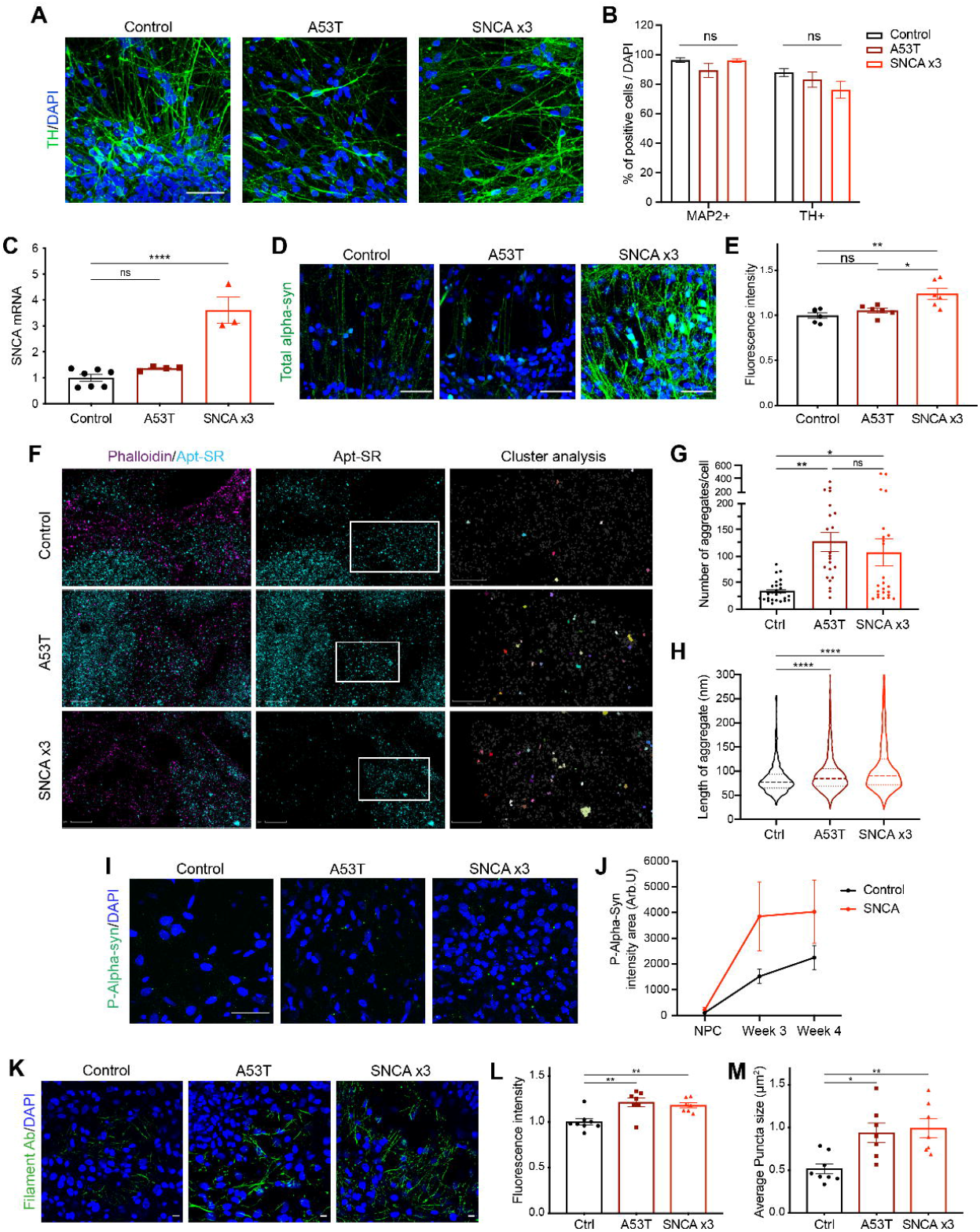
Generation of mDA neurons from hiPSC lines from patients with SNCA mutations display early alpha-synuclein aggregation. **(A)** Representative ICC images of control, A53T, and SNCA x3 patient hiPSCs derived mDA neurons showing high TH expression. Scale bar = 50μm. **(B)** Quantification of % MAP2 and TH positive cells after 3 weeks of differentiation (n = 2 control lines, 1 A53T lines, 1 SNCA x3 line, 1 neuronal induction, 5 fields of view per line, ns p > 0.05, two-way ANOVA). Values plotted as ±SEM. **(C)** Quantitative PCR showing relative mRNA expression of SNCA in mDA neurons at 3 weeks of differentiation (n = 3 control lines, 2 A53T lines, 1 SNCA x3 line, 3 neuronal inductions, ns p > 0.05, **** p < 0.0001, one-way ANOVA). Values plotted as ±SEM. **(D)** Representative ICC images showing the expression of alpha-synuclein at 4 weeks of differentiation (scale bar = 50μm). **(E)** Quantification of the relative normalised fluorescence intensity of alpha-synuclein at 4 weeks of differentiation (n = 2 control lines, 2 A53T lines, 1 SNCA x3 line, 1 neuronal induction, 3-6 fields of view per line, ns p > 0.05, * p < 0.05, ** p < 0.005, one-way ANOVA). Values plotted as ±SEM. **(F)** Super-resolved images from control, A53T, SNCA x3 1 week old mDA neurons. Left panel shows phalloidin and aptamer. Middle panel shows only superresolved aptamer binding events. Last panel shows a magnified version of the white box in middle panel showing the presence of aggregates picked up by DBSCAN. Left panel scale bar = 2 μm. Middle panel scale bar = 2 μm. Right panel = 1 μm. **(G)** Quantification showing the number of aggregates per cell in control, A53T and SNCA x3 mDA neurons (n = 5-9 cells, 2-3 fields of view per line, 2 control lines, 1 A53T line, 1 SNCA x3 line, ns p > 0.05, * p < 0.05, ** p < 0.005, one-way ANOVA). **(H)** Quantification showing the length of all aggregates in control, A53T and SNCA x3 mDA neurons represented in a violin plot (**** p < 0.0001, one-way ANOVA). **(I)** Representative ICC images showing phosphorylated alpha-synuclein in green at 4 weeks of differentiation in control, A53T, and SNCA x3 lines. Scale bar = 50μm. **(J)** Quantification showing the normalised phosphorylated alpha-synuclein area over time from NPCs through to 4 weeks of differentiation in control and SNCA PD lines (n = 2-3 fields of view per line at each time point, 2 control lines, 1 A53T line, 1 SNCA x3 line). **(K)** Representative ICC images showing the expression of filamentous aggregated forms of alpha-synuclein recognised by a conformation specific antibody, at 6 weeks of differentiation. Scale bar = 10μm. **(L)** Quantification of the normalised fluorescence intensity of aggregated filamentous forms of alpha-synuclein (n = 3 control lines, 2 A53T lines, 1 SNCA x3 line, 1 neuronal induction, 3-7 fields of view per line, ** p < 0.005, one-way ANOVA). Values plotted as ±SEM. **(M)** Quantification of the average puncta size of the aggregated filamentous forms of alpha-synuclein.

As validated in the control line mDA neurons, *SNCA* x3 neurons exhibited TTX-sensitive action potential generation in response to injected current, single channel NMDA receptor activity and stimulation evoked changes in capacitance (Supplementary Figure 5), confirming characteristic electrophysiological activity in patient-specific mDA neurons.

### Alpha-synuclein aggregation occurs early, and progresses over time in *SNCA* mDA neurons

During disease, monomeric α-synuclein aggregates to form small soluble oligomers, with accumulation of β-sheet structure, and later fibrils ^26,27^. We investigated whether *SNCA*-PD mDA neurons exhibited early aggregate formation using a single-stranded oligonucleotide aptamer, which binds to β-sheet rich protein structures containing α-synuclein ^28^. Due to the small size of oligomers (often below the diffraction limit of light ≈250 nm), we employed a previously described super-resolution technique, aptamer-based DNA-PAINT (point accumulation in nanoscale topography) to visualise and quantify α-synuclein oligomers. This consists of an aptamer probe that is conjugated to an oligonucleotide sequence which is recognised by the fluorophore-conjugated DNA imaging strand ^28^. After 1 week of differentiation, cells were incubated with the Aptamer, as well as the actin probe, Phalloidin-647. Using a combination of dSTORM (direct stochastic optical reconstruction microscopy) and DNA-PAINT, we superresolved oligomers and actin in mDA neurons to quantify aggregates based on a single-cell resolution (Figure 4F) (Supplementary Figure 6A). Performance of a clustering algorithm analysis, DBSCAN, quantified the clusters based on localisations that were within 60 nm of each other and had at least 15 localisations to be considered a cluster; to remove non-specific localisations (Last panel, Figure 4F). As early as 1 week of differentiation, prior to functional neuronal identity, both, A53T and *SNCA* x3 cells displayed a significantly higher number of aggregates per cell (Figure 4G) (Supplementary Figure 6B) (Ctrl = 33 ± 3.9, A53T = 127 ± 18.2, *SNCA* x3 = 106 ± 26.4, p < 0.05). In addition, the average length of the aggregates was higher in A53T, and *SNCA* x3 cells (Figure 4H) (Supplementary Figure 6C) (Ctrl = 83.4 nm ± 1.05, A53T = 94.9 nm ± 0.83, SNCA x3 = 113.0 nm ± 1.45, p < 0.0001). In order to rule out nonspecific binding of oligonucleotides, cells were incubated with only the imaging strand. The number of single-molecule localisations was low when incubated with only the imaging strand and was significantly increased upon addition of the aptamer (Supplementary Figure 6D-F), confirming that the majority of localisations detected are due to the aptamer binding to its target.

We next investigated whether aggregation worsens in neurons as they age. Phosphorylation of α-synuclein at Ser129 is a characteristic feature of Lewy body and Lewy neurite pathology in the post-mortem brain ^29^. Ser129-phosphorylated α-synuclein was detectable in early mDA neurons (3-4 weeks old), and increased with age (Figure 4I & J). As the neurons aged, both A53T and *SNCA* x3 neurons had a higher amount of phosphorylated α-synuclein compared to control neurons, suggesting that these neurons displaed characteristic aggregation phenotypes (Figure 4I & J). We also utilised a conformation-specific antibody that recognises all types of aggregated forms of α-synuclein ^30^. At 6 weeks of differentiation, we observed the presence of aggregated filamentous α-synuclein in both the A53T and the *SNCA* x3 lines which were significantly increased compared to control neurons (Figure 4K), confirmed by quantification of fluorescence intensity (Figure 4L) (Ctrl = 1.00 ± 0.04, A53T = 1.21 ± 0.05, *SNCA* x3 = 1.18 ± 0.03, p < 0.005) and the average size of the aggregate puncta (Figure 4M) (Ctrl = 0.52 μm^2^ ± 0.06, A53T = 0.94 μm^2^ ± 0.11, SNCA x3 = 0.99 μm^2^ ± 0.11, p < 0.05; p < 0.005).

Collectively, our data demonstrates that, using highly sensitive methods, we were able to detect and measure the formation of β-sheet rich, toxic, oligomeric species as the earliest pathological hallmark in *SNCA* neurons. Over time, aggregation progresses to form aggregates that become detectable by traditional immunocytochemical methods.

### Abnormal calcium signalling emerges early in PD patient-derived *SNCA* mDA neurons

Calcium dysregulation is one of the key phenotypes related to α-synuclein aggregates ^19,30^. We investigated whether physiological calcium responses were altered by the presence of *SNCA* mutations. We stimulated hiPSC-derived *SNCA* PD mDA neurons with KCl (50mM) to induce the opening of potential-sensitive Ca^2+^ channels in neurons at various stages of mDA differentiation. By two weeks into terminal differentiation, a similar proportion of control and *SNCA*-PD neurons (60-70%) showed a cytosolic response to KCl confirming the presence of voltage-gated calcium channels (Ctrl = 65.5% ± 5.7, A53T = 71.9% ± 1.7, SNCA x3 = 66%, p > 0.05) (Figure 5Ai & Aii), in a high neuronal culture (Ctrl = 90.8% ± 1.69, A53T = 90.55% ± 2.19, *SNCA* x3 = 91.6% ± 3.32) (Figure 5Aiii). However, neurons harbouring *SNCA* mutations displayed a significantly impaired recovery of cytosolic calcium in response to KCl from 2 weeks of differentiation (Ctrl = 1.00 ± 0.06, A53T = 0.22 ± 0.03, SNCA x3 = 0.19 ± 0.04, p < 0.0001) (Figure 5Aiv & Av).

**Figure 5.**
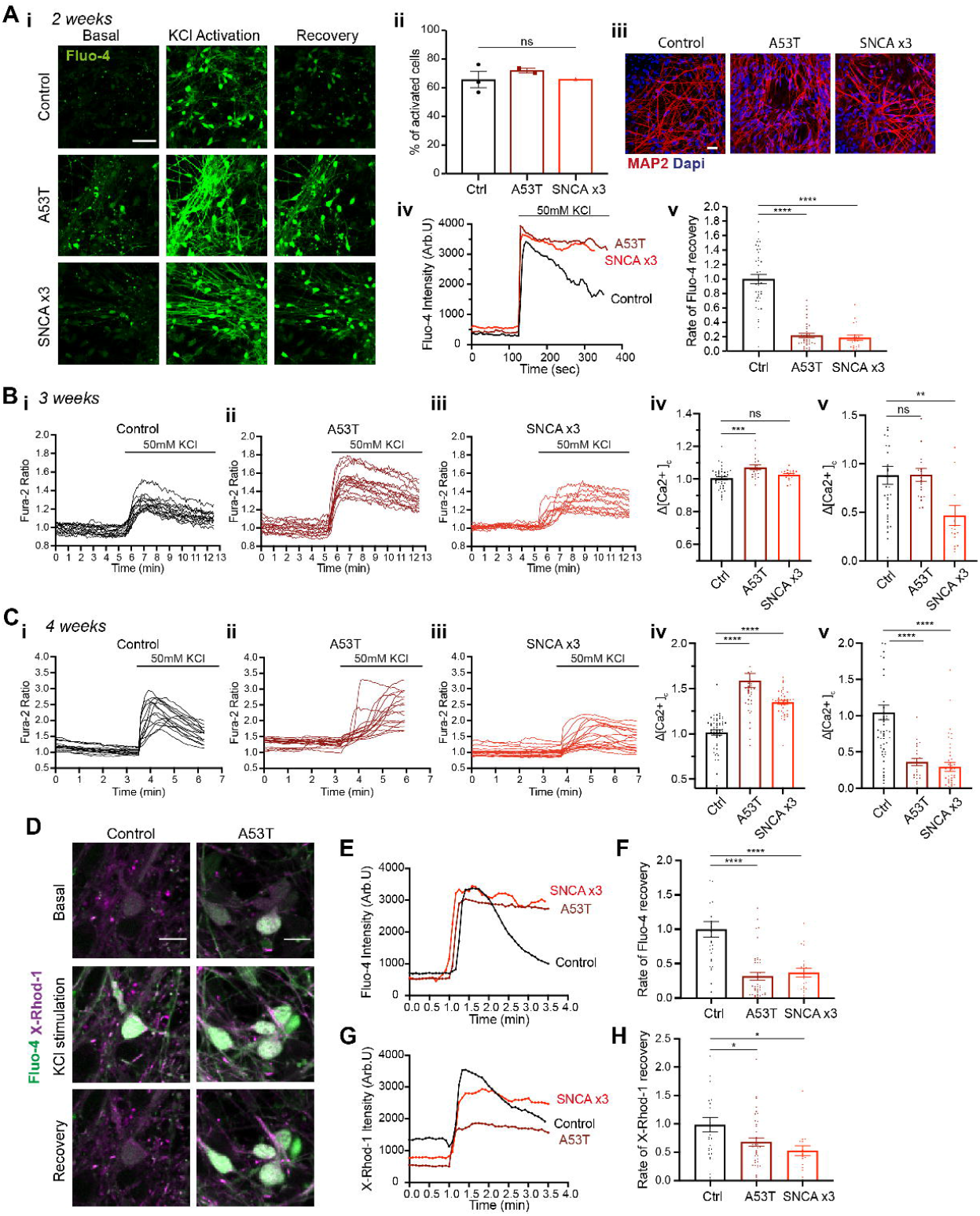
SNCA PD mDA neurons display early calcium dysregulation. **(A)** Neuronal characterisation at 2 weeks of differentation. **(i)** Representative time series Ca^2+^ images in response to KCl (scale bar = 50μm). **(ii)** Quantification of the number of cells with KCl induced calcium signal at 2 weeks of differentiation (n = 3 control hiPSC lines, 2 A53T patient lines, 1 SNCA x3 patient line, 1 neuronal induction, ns p > 0.05, one-way ANOVA). Values plotted as ±SEM. **(iii)** Representative images of neuronal marker expression **(iv)** Representative single-cell trace showing delayed recovery after KCl stimulation in patient mDA neurons. **(v)** Quantification of the normalised rate of recovery of Fluo-4 after stimulation with KCl (n = 15-20 cells per line, 1 neuronal induction, **** p < 0.0001, one-way ANOVA). **(B)** Representative traces showing the Fura-2 ratio in response to 50mM KCl in 3 week **(i)** control neurons, **(ii)** A53T neurons, and **(iii)** SNCA x3 neurons. **(iv)** Quantification of the basal calcium ratio (Δ[Ca^2+^]_c_) before KCl stimulation in 3 week old neurons (n = 15-20 cells per line, 2 control hiPSC lines, 1 A53T line, 1 SNCA x3 line, 1 neuronal induction, ns p > 0.05, *** p < 0.0005, one-way ANOVA). **(v)** Quantification of the rate of calcium (Δ[Ca^2+^]_c_) recovery in response to KCl in 3 week old neurons (n = 15-20 cells per line, ns p > 0.05, ** p = 0.0073, one-way ANOVA). Values plotted as ±SEM. **(C)** Representative traces showing the Fura-2 ratio in response to 50mM KCl in 4 weeks of differentiation **(i)** control neurons, **(ii)** A53T neurons, and **(iii)** SNCA x3 neurons. **(iv)** Quantification of the basal calcium ratio (Δ[Ca^2+^]_c_) before KCl stimulation in 4 week old neurons (n = 15-20 cells per line, 2 control lines, 1 A53T line, 1 SNCA x3 line, 1 neuronal induction, **** p < 0.0001, one-way ANOVA). **(v)** Quantification of the rate of calcium (Δ[Ca^2+^]_c_) recovery in response to KCl in 4 week old neurons (**** p < 0.0001, one-way ANOVA). Values plotted as ±SEM. **(D)** Representative time series snapshots of 4-5 week old control and A53T neurons loaded with Fluo-4 (green) and X-Rhod-1 (magenta) (scale bar = 10μm). **(E)** Representative single-cell trace showing delayed recovery of Fluo-4 after KCl stimulation in patient mDA neurons. **(F)** Quantification of the normalised rate of recovery of Fluo-4 after stimulation with KCl in 4-5 week old neurons (n = 2 control lines, 2 A53T lines, 1 SNCA x3 line, 1 neuronal induction, 10-20 cells per line, **** p < 0.0001, one-way ANOVA). Values plotted as ±SEM. **(G)** Representative single-cell trace showing delayed recovery of X-Rhod-1 after KCl stimulation in patient mDA neurons. **(H)** Quantification of the normalised rate of recovery of X-Rhod-1 after stimulation with KCl in 4-5 week old neurons (n = 10-20 cells per line, * p < 0.05, one-way ANOVA). Values plotted as ±SEM.

Using the ratiometric calcium indicator, Fura-2, we found that A53T mDA neurons displayed higher basal Δ[Ca^2+^]_c_ (Ctrl = 1.00 ± 0.01, A53T = 1.07 ± 0.01, *SNCA* x3 = 1.03 ± 0.01, p < 0.0005) (Figure 5Bi - iv) and calcium amplitude (Supplementary Figure 7A) than control neurons at 3 weeks post differentiation. In addition, we found a delayed calcium recovery rate in *SNCA* x3 mDA neurons by KCl stimulation which indicates an impaired cytosolic calcium efflux (Ctrl = 0.88 ± 0.09, A53T = 0.89 ± 0.07, *SNCA* x3 = 0.47 ± 0.10, p = 0.0073) (Figure 5Bi – iii & Bv). After 4 weeks of differentiation, both A53T and *SNCA* x3 mDA neurons displayed a higher basal Δ[Ca^2+^]_c_ (Ctrl = 1.02 ± 0.03, A53T = 1.59 ± 0.08, *SNCA* x3 = 1.35 ± 0.02, p < 0.0001), as well as a delayed recovey following KCl stimulation (Ctrl = 1.04 ± 0.1, A53T = 0.36 ± 0.05, *SNCA* x3 = 0.29 ± 0.06, p < 0.0001) compared to control neurons (Figure 5C). In addition, *SNCA* x3 neurons had a lower calcium amplitude compared to control neurons (Supplementary Figure 7A).

During cytosolic calcium influx, mitochondria act as the major buffer for calcium, shaping the cytosolic calcium signal and maintaining calcium homeostasis ^31^. Upon KCl-induced calcium influx, we observed a concomitant increase in mitochondrial calcium, measured by the fluorescent dye X-Rhod-1 (Figure 5D). However, the recovery of the mitochondrial calcium signal was significantly delayed in the *SNCA*-mutant neurons compared to controls, suggesting a potential impairment of mitochondrial calcium efflux (Figure 5E – H) (X-Rhod-1: Ctrl = 0.98 ± 0.13, A53T = 0.68 ± 0.07, SNCA x3 = 0.53 ± 0.09, p < 0.05) (Fluo-4: Ctrl; = 1.00 ± 0.11, A53T = 0.32 ± 0.05, SNCA x3 = 0.37 ± 0.06, p < 0.0001). Taken together, these results suggest calcium dysregulation is the first functional phenotype to emerge, early after differentiation.

Oxidative stress, lysosomal dysfunction, and alterations in mitochondria emerge later in PD patient derived-*SNCA* mDA neurons. We assessed mitochondrial function of *SNCA*-mutant PD neurons using the lipophilic cationic dye TMRM, which accumulates less in depolarised mitochondria upon the loss of mitochondrial membrane potential. Depolarisation of the mitochondria emerges at 4 weeks of differentation in *SNCA*-mutant mDA neurons shown by a reduction in mitochondrial membrane potential (Figure 6A & B) (Ctrl = 100% ± 3.4, A53T = 74.1% ± 3.3, *SNCA* x3 = 80.0% ± 7.1, p < 0.05; p < 0.005) (Supplementary Figure 7C). Changes in the mitochondrial network are also apparent in *SNCA*-mutant neurons, with a reduced area/volume occupied by mitochondria (iso-Ctrl of *SNCA* x3 = 100% ± 6.8, *SNCA* x3 = 74.2% ± 7.5, p < 005.; iso-Ctrl of A53T = 100% ± 7.4, A53T = 74.4% ±4.7, p < 0.005) (Supplementary Figure 7D), which was reversed in isogenic controls.

**Figure 6.**
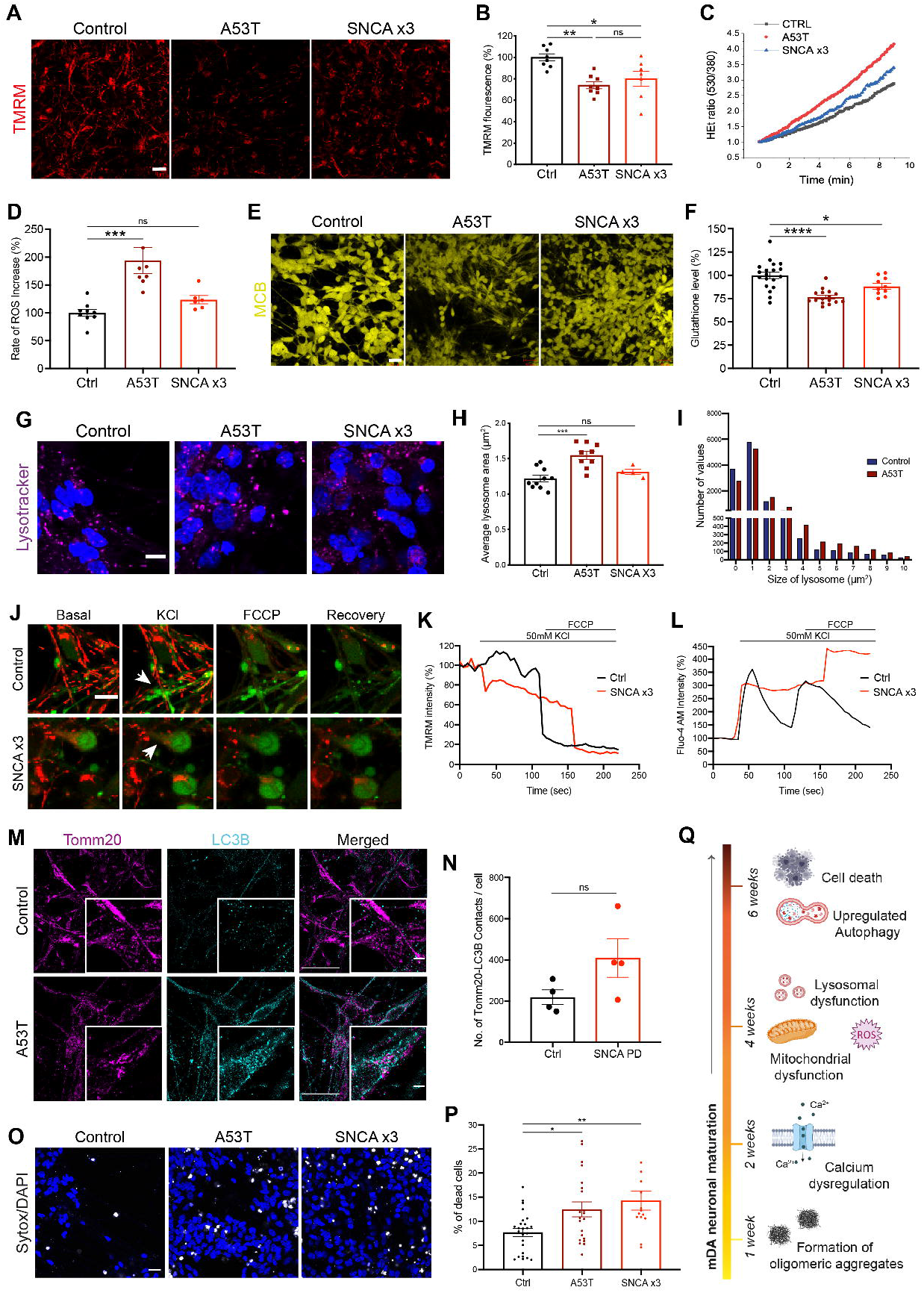
Cellular dysfunction and cell death arise later in SNCA PD mDA neurons. **(A)** Representative live-cell imaging of mitochondrial fluorescence using the lipophilic cationic dye TMRM at 4 weeks of differentiation. Scale bar = 10μm. **(B)** Quantification of the normalised fluorescence intensity of TMRM (n = 3-4 fields of view per line, 2 coverslips per line, across 2 control lines, 2 A53T lines, and 1 SNCA x3 line, 2 neuronal inductions, ns p > 0.05, * p < 0.05, ** p < 0.005, one-way ANOVA). Values plotted as ±SEM. **(C**) Trace showing the ratiometric measurement of superoxide generation using dihydroethidium (HEt) at 4 weeks of differentiation. **(D)** Quantification of the rate of superoxide generation based on HEt ratiometric fluorescence (n = 1-2 coverslips per line, across 3 control lines, 2 A53T lines, and 1 SNCA x3 line, 3 neuronal inductions, ns p > 0.05, *** p < 0.0005, one-way ANOVA). Values plotted as ±SEM. **(E)** Representative live-cell imaging of endogenous glutathione using the fluorescent reporter MCB at 4 weeks of differentiation. Scale bar = 20μm. **(F)** Quantification of the endogenous level of glutathione based on MCB fluorescence (n = 4-6 fields of view per line, 3 control lines, 2 A53T lines, and 1 SNCA x3 line, 2 neuronal inductions, * p < 0.05, **** p < 0.0001, one-way ANOVA). Values plotted as ±SEM. **(G)** Representative live-cell imaging of lysosomes and nuclear marker Hoechst 33342 at 4 weeks of neuronal differentiation (scale bar = 5μm). **(H)** Quantification of the average lysosomal area after 4 weeks of differentiation (n = 4-6 images per line, across 2 control lines, 2 A53T lines, and 1 SNCA x3 line, 1 neuronal induction ns p > 0.05, *** p < 0.0005, one-way ANOVA). Values plotted as ±SEM. **(I)** Histogram plot showing the relative number of lysosomes in each set area bin (0-10μm^2^) (n = 3-5 images per line, across 2 control lines, 2 A53T lines, 1 neuronal induction). **(J)** Representative single-cell trace showing TMRM intensity in response to KCl and FCCP in control and SNCA x3 6 week old neurons (n = 5-10 cells per line, 1 control line, 1 A53T line, 1 SNCA x3 line, 1 neuronal induction). **(K)** Representative single-cell trace showing Fluo-4 intensity in response to KCl and FCCP in control and SNCA x3 6 week old neurons. **(L)** Representative time series snapshots of TMRM (red) and Fluo-4 (green) in 6-week old neurons showing the response to KCl and FCCP. Arrow in the control cell highlights polarised mitochondria and calcium response to KCl and FCCP. Arrowhead in the SNCA x3 cell highlights KCL induced TMRM intensity decrease (scale bar = 10μm). **(M)** Structured illumination microscopy (SIM) images of control and A53T cells probed for mitochondrial marker Tomm20, and the autophagosome marker LC3B, in 5 week terminally differentiated mDA neurons (scale bar = 20μm). Smaller image depicts a zoom showing morphology and colocalization of the markers (scale bar = 2μm). **(N)** Quantification of the number of Tomm20-LC3B colocalizations per cell (n = 1-2 images per line, 2 control lines, 2 A53T lines, 1 SNCA x3 line, 1 neuronal induction, ns p > 0.05 (p = 0.11), unpaired t-test). Values plotted as ±SEM. **(O)** Live cell images depicting dead cells in mDA neurons at 4 weeks of terminal differentiation using the fluorescent dye SYTOX green (scale bar = 20μm). **(P)** Quantification of the percentage of dead cells (n = 4-5 fields of view, 1-2 wells per line, 2 control lines, 2 A53T lines and 1 SNCA x3 line, 2 neuronal inductions, * p < 0.05, ** p < 0.005, one-way ANOVA). **(Q)** A schematic illustration showing temporal sequence of cellular phenotypes in human PD model. The earliest abnormality is the accumulation of small aggregates with a specific beta sheet conformation prior to the development of molecular and functional identiy of mDA neurons (at 1 week of differentation). When the midbrain neurons exhibit functional specialisation by 2 weeks of differentiation, impaired calcium signalling is displayed followed by mitochondrial dysfunction and oxidative stress upon confimraiton fo midbrain identity by 4 weeks of differentation. As the maturation is progressed upregulated autophagy and cell death are induced (at 6 weeks of differentation).

Mitochondrial dysfunction can result in overproduction of mitochondrial and extra-mitochondrial reactive oxygen species (ROS). We measured the generation of cytosolic ROS using the fluorescent reporter dihydroethidium (HEt), which changes fluorescence emission upon oxidation. The rate of superoxide production was significantly higher in the A53T lines, and approximately 20% higher in the *SNCA* x3 line compared to controls (Ctrl = 100% ± 6.3, A53T = 194.0% ± 23.7, SNCA x3 = 123.6% ± 7.5, p < 0.0005) (Figure 6C & D, Supplementary Figure 7B). We also measured the levels of the endogenous antioxidant glutathione using the Monochlorobimane (MCB) fluorescence indicator, which showed a significant reduction of glutathione in both, A53T and *SNCA* x3 lines compared to the controls (Ctrl = 100% ± 3.5, A53T = 76.7% ± 1.9, SNCA x3 = 88.2% ± 3.1, p < 0.0001) (Figure 6E & F, Supplementary Figure 7E). The combination of increased ROS production together with depleted antioxidant levels suggests that both, A53T and *SNCA* x3 mDA neurons are under oxidative stress.

We measured the lysosomal compartment in *SNCA*-PD neurons using LysoTracker™ Deep red; a cationic fluorescent dye that only accumulates in acidic cellular compartments (lysosomes). We found A53T neurons from 4 weeks differentation exhibited significantly larger lysosomes compared to the healthy control neurons, with a notable increase in lysosomal area in A53T neurons compared to controls (Ctrl = 1.2 μm^2^ ± 0.05, A53T = 1.5 μm^2^ ± 0.06, *SNCA* x3 = 1.3 μm^2^ ± 0.04, p < 0.0005) (Figure 6G & H). There was also an increase in the number of lysosomal spots in both A53T and *SNCA* x3 mDA neurons, which is reversed in isogenic controls (iso-Ctrl of *SNCA* x3 = 100% ± 6.6, *SNCA* x3 = 73.5% ± 8.7, p < 0.05; iso-Ctrl of A53T = 100% ± 6.4, A53T= 78.4% ±4.6, p < 0.005) (Supplementary Figure 7F). Control neurons had a higher proportion of smaller lysosomes (mainly between 0-2 μm^2^) compared to the A53T neurons, which had a higher proportion of larger lysosomes (mostly 2-10 μm^2^) (Figure 6I), suggesting that the lysosomes are swollen, implying lysosomal dysfunction.

### Mitochondrial fragmentation, autophagy, and cell death occur in aged PD patient-derived *SNCA* mDA neurons

To test the consequence of aberrant calcium flux and mitochondrial dysfunction, we loaded *SNCA* and control neurons with cytosolic and mitochondrial calcium and membrane potential indicators and tested their response to KCl. As expected, KCl induced opening of the voltage-dependent calcium channels, and a cytosolic calcium signal. This was associated with mitochondrial influx of calcium, during which the mitochondria retained their mitochondrial membrane potential (Figure 6J-L). *SNCA* mutant mDA neuron mitochondria exhibited depolarisation upon calcium influx into the mitochondria, which was induced by the cytosolic calcium signal (Figure 6J & K, Supplementary Figure 7G), concurrently showing an impaired recovery of cytoplasmic calcium (Figure 6L). Rapid depolarisation events in the mitochondria may reflect transient permeability transition pore (PTP) opening, an event used for mitochondrial efflux during mitochondrial calcium overload.

Damaged organelles, including mitochondria, as well as protein aggregates are cleared through the autophagy-lysosomal pathway. Autophagy results in the formation of autophagosomes, which engulf the target, and process them to lysosomes for destruction. As we found both aggregation and mitochondrial damage present in *SNCA* mutant PD mDA neurons, we studied the expression of the autophagosome marker LC3B together with the mitochondrial marker Tomm20 at week 5 post differentiation using a super-resolution approach, structured illumination microscopy (SIM) to resolve the contacts between the mitochondria and the autophagosomes. Control cells exhibited Tomm20 staining with an intact mitochondrial network and exhibited a low level of LC3B staining (top panels, Figure 6M). In contrast, the Tomm20 appearance in the A53T lines exhibited fragmented mitochondria with a disrupted mitochondrial network, and there was an increase in the LC3B staining suggesting an autophagosome response (lower panels, Figure 6M). We also observed an increase in the number of colocalisations between Tomm20 and LC3B in all *SNCA*-PD lines compared to control lines, suggesting close contact between autophagosomes and the mitochondria (Ctrl = 219 ± 36, SNCA PD = 410 ± 94, Tomm20-LC3B colocalizations per cell, p > 0.05) (Figure 6N).

Finally, we examined the basal viability of the *SNCA* mutant neurons compared to the controls using the fluorescent dye SYTOX™ Green to identify dead cells. We observed a significant increase in cell death in both the A53T and *SNCA* x3 lines compared to the healthy control lines at 4-5 weeks post differentiation (Ctrl = 7.7% ± 0.8, A53T = 12.5% ± 1.5, SNCA x3 = 14.3% ± 2.0, p < 0.05) (Figure 6O & P) as well as their isogenic pair control neurons (iso-Ctrl of *SNCA* x3 = 100% ± 27.3, *SNCA* x3 = 247.3% ± 35.9,p < 0.005 = iso-Ctrl of A53T = 100% ± 10.5, A53T = 285.3% ± 31.1, p < 0.0005) (Supplementary Figure 7H), suggesting that aggregate formation, impaired calcium, protein and mitochondrial homeostasis is toxic to mDA *SNCA*-PD neurons (Figure 6Q).

## Discussion

Lineage restriction to diverse cellular fates in the neuraxis is the consequence of interplay of multiple developmental signals, which are regulated in a spatio-temporal manner. *In vitro*, such lineage restriction can be achieved using a small molecule based approach. In keeping with previous protocols, we initiated neuronal induction via small molecule dual-SMAD inhibition ^6^, caudalisation to the midbrain via timed activation of Wnt signalling through a GSK-3β inhibitor, and floor-plate ventralization through Shh signalling activated by small molecule agonism. This yielded a population of enriched mDA progenitors, which has been shown to be vitally important in mouse graft outcome ^32,33^.

Enriched mDA progenitors were subsequently differentiated into enriched mDA TH-positive neurons (>80%) after 3 weeks, using a combination of two molecules (a ROCK inhibitor and a Notch inhibitor) to allow the mDA NPCs to survive and exit the cell cycle, promoting differentiation into post-mitotic mDA neurons. Diverging from established methods, the mDA NPCs successfully differentiated into enriched mDA neurons without the use of any neurotrophic growth factors ^32^, improving the cost-effectiveness of this approach. Single-cell sequencing identified seven neuronal clusters, which expressed key markers of mDA neurons, and a further five clusters with key markers of midbrain NPCs. RNA-velocity and latent time analyses ^20,34^ revealed the developmental trajectories within the culture, with NPC clusters representing early cell types that were predicted to generate mDA clusters, of varying identities. Furthermore, we identified ‘driver genes’ that were associated with these trajectories, highlighting a potential role for *STMN2, DCX*, and *SYT1* in midbrain neuronal development. ^35,36^. Our mDA neurons displayed functional properties including calcium channel activity, functional DAT activity, and the synthesis, metabolism and secretion of dopamine. The mDA neurons also display typical neuronal electrophysiological behaviour, pre- and postsynaptic currents, and mature into functional neuronal networks. Differentiating hiPSCs from patients with PD yielded highly enriched mDA neurons of similar molecular and functional identity as control hiPSC-derived mDA neurons, confirming that this differentiation approach may be used to generate models of disease.

*In vivo* and *in vitro* models of PD have revealed several putative mechanisms that may cause neuronal toxicity in disease, but virtually all models exhibit several forms of cellular stress occurring simultaneously. We harnessed the developmental nature of the hiPSC system to assess the temporal sequence of events that unfold in the context of *SNCA* mutations, in order to distinguish early, and likely causative events in the cell, from late bystander events. The critical hallmark of the synucleinopathies, and therefore the key phenotype for a model to recapitulate, is the detection of aggregated forms of α-synuclein in neurons. Aggregate load in A53T patient lines are reported to be increased due to the mutation ^37^, and in SNCA x3 patient lines, are higher due to an increase in cellular α-synuclein ^38^. Aggregates have been reported at day 35 in mDA cultures ^39^. We have previously used single molecule and super-resolution approaches ^13^, to detect the formation of the earliest small soluble oligomers outside cells and within cells, and we defined the most toxic aggregate species as a small soluble oligomer with cross β-sheet structure ^26,30,40^. Here, we applied highly sensitive approaches (super-resolution microscopy) to the *SNCA* mDA neurons to define when oligomer aggregation starts. Small oligomeric species of α-synuclein were detected early into differentiation (1 week), prior to the molecular and functional specification of midbrain dopaminergic neurons, but after the expression of *SNCA* increased from NPC stage. Both the number and size of these small oligomeric aggregates are higher in *SNCA* A53T and *SNCA* x3 PD lines, and, furthermore, a specific conformation of α-synuclein, that is, the β-sheet-rich α-synuclein oligomers, occurs at early stages. These small oligomers are highly hydrophobic and their accumulation is likely to be responsible for the toxicity of cells ^27,41,42^. Abnormal aggregation of α-synuclein was a persistent phenomenon, and phosphorylated α-synuclein puncta were apparent using diffraction limited microscopy at 3 weeks, and the accumulation of phosphorylated forms progressed over time in culture. Finally, conformation-specific antibodies detected fibrillar forms of α-synuclein at later time points, to week 6 of differentiation.

The hydrophobic nature of the oligomer is known to induce a range of cellular stresses, due to its ability to insert and disrupt membranes ^43,44^. Notably, the earliest functional phenotype observed in this model, is calcium dysregulation at 2-3 weeks of differentiation. We noted higher basal levels of cytosolic calcium, an increased calcium influx on stimulation, and a delayed calcium recovery in *SNCA*-PD mDA neurons. We have previously shown that the β-sheet-rich oligomer is able to permeabilise cell membranes due to its lipophilic properties, and induce calcium fluxes, leading to increased cytosolic calcium in response to glutamate and KCl ^19,30^. Calcium dysregulation has also been reported in other synucleinopathy models, where it is reported that oligomers interact with receptors and calcium channels ^45,46^.

Mitochondria are fundamental for cellular function and homeostasis, in particular for buffering cytosolic calcium, and ATP generation, which is a large source of ROS generation ^31^. Impairment in calcium fluxes plays an important role in mitochondrial oxidant stress ^47^, highlighting the delicate balance between mitochondrial and cellular calcium homeostasis. We previously showed that oligomeric structures of α-synuclein interact with ATP synthase, causing mitochondrial dysfunction and early opening of the mitochondrial PTP, which causes neuronal death in *SNCA* x3 neurons ^13^. Similarly, oligomers have been shown to induce complex I-dependent mitochondrial dysfunction through mitochondrial calcium, which induced swelling and cytochrome C release ^48^. A53T hiPSC-derived neurons exhibit an increase in nitrostative, and ER stress ^16^, as well as mitochondrial dysfunction ^39^. Oligomers further induce the generation of aberrant ROS in both the cytosol and mitochondria ^13,18,30^. In this study, we observed abnormalities in mitochondrial function including a reduced membrane potential, abnormal mitochondrial calcium efflux, and fragmentation of the mitochondrial network. Cytosolic calcium fluxes induced rapid mitochondrial membrane depolarisation, reflecting early PTP opening induced by mitochondrial calcium overload. Oxidative stress was evident, based on aberrant generation of superoxide, with a concomitant reduction in glutathione, at 4 weeks post differentiation.

Autophagy maintains cellular function through lysosome-dependent degradation of damaged organelles or aggregates ^49^. In addition to mitochondrial and oxidative stress at a similar time point, we detected a swelling of the lysosomes in the *SNCA* A53T lines. α-synuclein-dependent impairment of lysosomal capacity has been previously reported ^17^. In addition to lysosomal alterations, super-resolution microscopy revealed *SNCA*-PD mDA neurons exhibit an increase in autophagosomes, that are in close physical contact with fragmented mitochondria, similar to previous studies ^50^. Our results support a mechanistic hypothesis that in the disease process, abnormal α-synuclein leads to three major effects in the cell: (i) β-sheet-rich oligomeric species disrupt cellular membranes resulting in early cytosolic calcium phenotypes, (ii) oligomers induce disruption of mitochondrial function, oxidative stress and fragmentation of the mitochondrial network, and (iii) a lysosomal response to potentially clear the misfolded α-synuclein. Later, the accumulation of both damaged mitochondria and misfolded protein stimulates an autophagic response in the *SNCA* models.

In summary, we successfully generate highly enriched population of mDA neurons from hiPSCs, that express mDA markers, functional dopamine transport and form neuronal networks. By enriching the cell of interest in patient-derived lines with *SNCA* mutations, we were able to delineate that the earliest abnormality was the accumulation of small hydrophobic aggregates with a specific β-sheet conformation. These aggregates induce early calcium dysregulation, and this is followed by mitochondrial dysfunction and oxidative stress, and later by compensatory upregulation of autophagy. Dissecting the temporal sequence of pathological events revealed the first and critical driver of pathogenesis, and subsequent organellar impairment, is the development of toxic protein aggregates.

## Acknowledgements

We would wish to thank the patients for the fibroblast donation. We would also like to thank the Francis Crick Institute Flow Cytometry, Advanced Light Microscopy, Advanced Sequencing, and Bioinformatics and Biostatistics STPs for their help and equipment in conducting and analysing the flow cytometry, fluorescence microscopy, and single-cell RNA-seq experiments.

## Author Contributions

Conceptualization, G.S.V, R.P, and S.G.; Methodology, S.G, R.P, M.H.H, and A.Y.A.; Investigation, G.S.V, M.L.C, J.R.E, Z.Y, A.I.W, D.M, J.P.L, and P.R.A. RNA-seq, S.S.; HPLC, H.A, S.E, and S.H.; Electrophysiology, S.S.; Writing-original draft, G.S.V.; Writing-review & Editing, S.G, R.P, and D.A.; Funding acquisition, S.G.

## Methods

### Data availability

The data that supports the findings of this study is available on the open access repository Zenodo DOI: https://doi.org/10.5281/zenodo.7138359. Single-cell RNA-seq raw data were deposited to NCBI Gene Expression Omnibus. The accession code for the data is: GSE213569 (https://www.ncbi.nlm.nih.gov/geo/). The code used for all scRNA-seq and RNA velocity has been deposited online (DOI: https://doi.org/10.5281/zenodo.7260558), (https://github.com/strohstern/Transcriptomic_signatures_iPSC_derived_dopamine_neurons). Protocols used in this study can be found on the repository Protocols.io and the DOIs can be found in Supplementary table 5.

### Human induced pluripotent stem cell culture

Human induced pluripotent stem cells (hiPSCs) were maintained in feeder-free monolayers on Geltrex (ThermoFisherScientific) and fed daily with Essential 8 medium (Life Technologies). When confluent, hiPSCs were passaged using 0.5 μM EDTA (Life Technologies). All cells were maintained at 37°C and 5% carbon dioxide. The Isogenic control line of *SNCA* A53T was generated using CRISPR/Cas9 editing by Applied StemCell *Inc*. (USA, project ID: C1729). The isogenic control line of SNCA x3 was kindly provided from the Tilo Kunath lab which was also generated using CRISPR/Cas9 editing as published ^51^, Centre for Regenerative Medicine, Institute for Stem Cell Research, School of Biological Sciences, The University of Edinburgh, UK. All hiPSC lines used in this study are described in Supplementary table 1.

### Midbrain dopaminergic neuron (mDA) differentiation

For all mDA differentiations, hiPSCs were grown to 100% confluency. Differentiation was triggered by removing old media and replacing it with a chemically defined medium consisting of DMEM/F12, N2 supplement, Neurobasal, B27 supplement, L-Glutamine, non-essential amino acids, 50U/ml penicillin-streptomycin, β-mercaptoethanol (all from ThermoFisherScientific) and 5μg/ml insulin (Sigma), termed “N2B27”. Cells were patterned for 14 days with daily media changes. For the first 2 days, the media was supplemented with the small molecules 5μM SB431542 (Tocris Bioscience), 2μM Dorsomorphin (Tocris Bioscience), 1μM CHIR99021 (Miltenyi Biotec). On day 2, 1μM Purmorphamine (Merck Millipore) was added. On day 8, CHIR99021, and SB431542 were removed leaving only Dorsomorphin and Purmorphamine in the medium until day 14. Cells were enzymatically dissociated and split on days 4, 10, and 14 using 1mg/ml of Dispase (ThermoFisherScientific). After patterning, mDA neuronal precursor cells (NPCs) were maintained in N2B27 for 4 days. On day 19, cells were plated onto Geltrex pre-coated Ibidi 8-well chambers (100k/well), clear bottom 96-well plates (50k/well), or 12-well plates (500k/well), and terminally differentiated using N2B27 supplemented with 0.1μM Compound E (Enzo Life Sciences) and 10μM Y-27632 dihydrochloride (Rho kinase ROCK inhibitor) (Tocris) from day 20 for the whole duration of terminal differentiation, with two weekly media changes. NPCs could be cryopreserved on day 19, before terminal differentiation. For super-resolution microscopy, cells were plated on glass-bottom Ibidi 8-well chambers which were pre-coated with poly-D-lysine (PDL) overnight, followed by laminin (Sigma, L2020) in PBS for 1 hour.

### Immunocytochemistry (ICC)

For ICC, cells at the desired time point of differentiation had the media removed, followed by one wash in PBS, and fixed with 4% paraformaldehyde for 15 minutes at room temperature (RT). The paraformaldehyde was then removed, and cells were washed once in PBS. For the LC3B antibody (Cell Signalling Technologies #3868), samples were incubated in −20°C methanol after paraformaldehyde fixation. All samples were then blocked for non-specific binding and permeabilized in 5% bovine serum albumin (BSA) (Sigma) + 0.2% Triton X-100 (Sigma) in PBS for 60 minutes. The primary antibodies were then made up to the desired dilution (Supplementary Table 2) in 5% BSA and applied to the cells overnight at 4°C. The cells were then washed twice in PBS followed by the application of species-specific, secondary antibodies conjugated to relative Alexa Fluor dyes (ThermoFisherScientific), at a 1:500 dilution made up in 5% BSA to cells for 60 minutes at RT in the dark. After secondary antibody incubation, cells were washed once in PBS before being stained with 4’,6-diamidino-2-phenylindole nuclear stain (DAPI) in PBS for 5 minutes at a 1:1000 dilution. After DAPI incubation, cells were washed once with PBS before being submerged in fluorescence mounting medium (Dako), and stored at 4°C until imaging.

The samples were imaged using Zeiss 880 confocal system with a 40x, 1.4 N.A. oil objective, and a pinhole of 1 airy units (AU). Between 3-5 images were collected per sample, all with a Z projection consisting of 5 slices, and displayed as a maximum projection. Samples were also imaged using the PerkinElmer Opera Phenix™ High Content Screening System with 20 and 40x water objective lenses. A minimum of 5 fields of view and a Z projection of 3 slices was acquired per well, with the images displayed as a maximum projection. The accompanying software, Columbus™ was used to store and analyse acquired images (https://biii.eu/columbus-image-data-storage-and-analysis-system). The settings for the acquisition of images were kept the same for all samples in the experiment set.

### RNA extraction and quantitative polymerase chain reaction (qPCR)

RNA was harvested from snap frozen cell pellets using the Maxwell®ū RSC simplyRNA Cells kit (Promega), and the accompanying Maxwell®□ RSC 48 instrument. After RNA extraction, the RNA concentration and quality using the 260/280 ratio was assessed using the nanodrop. Up to 1μg of RNA was retro-transcribed into cDNA using the High-Capacity cDNA Reverse Transcription kit (ThermoFisherScientific). The qPCR was performed using TaqMan^™^ Gene Expression Assay (ThermoFisherScientific). For each gene, TaqMan^™^ probes were used (Supplementary Table 3) along with the TaqMan^™^ master mix, and sample cDNA following the manufacturer’s protocol. Samples, along with a minus reverse transcriptase control (-RT) were ran for each gene on the QuantSudio 6 Flex Real-Time PCR System (Applied Biosystems). The -RT served as a negative control, and the gene expression levels were normalised to the housekeeping gene GAPDH following the delta-delta Ct method. Gene expression values were expressed as the normalisation to either hiPSCs or mDA NPCs.

### Single-cell RNA-seq

#### Single cell generation, cDNA synthesis, library construction, and sequencing protocol

After 4 weeks of differentiation, mDA neurons from 3 control hiPSC lines were washed with PBS once, and then incubated with Accutase (Gibco^™^) for 5 minutes to obtain a single cell suspension. The samples were then diluted 1/3 before usage. The quality and concentration of each single-cell suspension was measured using Trypan blue and the Eve automatic cell counter. Each sample presented a concentration between a 1200-1700 cell/μl and viability ranged between 55-68%, samples with a viability above 57% were used for sequencing. Approximately 10000 cells were loaded for each sample into a separate channel of a Chromium Chip G for use in the 10X Chromium Controller (cat: PN-1000120). The cells were partitioned into nanoliter scale Gel Beads in emulsions (GEMs) and lysed using the 10x Genomics Single Cell 3’ Chip V3.1 GEM, Library and Gel Bead Kit (cat: PN-1000121). cDNA synthesis and library construction were performed as per the manufacturer’s instructions. The RNA was reversed transcribed and amplified using 12 cycles of PCR. Libraries were prepared from 10μl of the cDNA and 13 cycles of amplification. Each library was prepared using Single Index Kit T Set A (cat: PN-1000213) and sequenced on the HiSeq4000 system (Illumina) using 100 bp paired-end run at a depth of 65-100 million reads. Libraries were generated in independent runs for the different samples.

#### Pre-processing single-cell RNA-seq (scRNA-seq) data

Using the Cell Ranger (RRID:SCR_017344, https://support.10xgenomics.com/single-cell-gene-expression/software/pipelines/latest/what-is-cell-ranger) v3.0.2 Single-Cell Software Suite from 10X Genomics reads were aligned to the human reference genome (Ensembl release 93, GRCh38) (https://support.10xgenomics.com/single-cell-gene-expression/software/pipelines/latest/what-is-cell-ranger). The analysis was carried out using Seurat (RRID:SCR_016341) v3.1.0 ^52,53^ (https://satijalab.org/seurat/get_started.html) in R-3.6.1 (R Core Team, 2019 (RRID:SCR_001905, http://www.r-project.org/)). The code used for analysis has been deposited online (https://github.com/strohstern/Transcriptomic_signatures_iPSC_derived_dopamine_neurons).

Cells expressing fewer than 200 genes were excluded from the subsequent analysis. Using default parameter within Seurat v3.1.0 ^52,53^ data for each sample were normalised across cells using the ‘LogNormalize’ function with a scale factor of 10,000. A set of highly variable genes was identified using the ‘FindVariableFeatures()’ function (selection.method = “vst”, nfeatures = 2000). Data were centred and scaled using the ‘ScaleData()’ function with default parameters. Using the highly variable genes, PCA was performed on the scaled data and the first 30 principal components were used to create a Shared Nearest Neighbour (SNN) graph using the ‘FindNeighbors()’ function (k.param = 20). This was used to find clusters of cells showing similar expression using the ‘FindClusters()’ function across a range of clustering resolutions (0.2-1.4 in 0.2 increments). Based on the visualisation of average mitochondrial gene expression across different cluster resolutions using the R package Clustree v0.4.1 (RRID:SCR_016293) ^54^ https://CRAN.R-project.org/package=clustree) we selected a clustering resolution of 1.0 to exclude cluster with an average mitochondrial gene expression above 7.5% and, concomitantly, an average number of detected features below 1300.

#### Integration across samples

After filtering of cells/clusters based on mitochondrial gene expression and the number of detected features, we integrated the three samples using the standard workflow from the Seurat v3.1.0 package ^53^. After data normalisation and variable feature detection in the individual samples (see above), anchors were identified using the ‘FindIntegrationAnchors()’ function and datasets were integrated with the ‘IntegrateData()’ across 50 dimensions for all detected features in the datasets. We then performed dimension reduction (PC1-45) and cluster identification at resolutions 1.4. After removal of a cluster consisting mostly of suspected doublet cells identified using DoubletFinder ^55^, we performed data scaling including cell cycle score regression, dimension reduction (PC1-50) and cluster identification (resolutions 0.4-1.8).

Biomarker of each cluster were identified using Seurat’s ‘FindAllMarkers()’ function using the Wilcoxon rank sum test. We limited the test to positive markers for each cluster in comparison to all remaining cells. The positive marker genes had to be detected in 25% of cells in either of the two groups, with limiting testing further to genes which show, on average, at least 0.25-fold difference (log-scale) between the two groups of cells. Cluster identity was determined using visual inspection focusing on the expression of known marker genes.

#### RNA velocity estimation

For RNA velocity analysis, the spliced and unspliced reads were counted with alevin ^56^ as recommended ^57^ The count matrices were added to the pre-existing Seurat object which was subsequently used as input into scVelo (v0.2.2, RRID:SCR_018168, https://github.com/theislab/scvelo) to calculate RNA velocity values for each gene of each cell. scVelo was used in the “dynamical” mode with default settings. The resulting RNA velocity vector was embedded into the PCA and UMAP space by translating the RNA velocities into likely cell transitions using cosine correlation to compute the probabilities of one cell transitioning into another cell. We identified driver genes, i.e. those genes that show dynamic behaviour, as those genes with a fit likelihood in the dynamical model > 0.3. We also used PAGA ^34^ to perform trajectory inference for which directionality was inferred from the RNA velocities.

### Flow Cytometry

The protocol for staining cells for flow cytometry analysis was adapted from a previous study ^58^. Cells were washed once with PBS, before being detached into a single cell suspension using Accutase (Gibco^™^). A cell suspension of 500k/ml was prepared in media. Cells were then centrifuged at 200g for 5 minutes, and the supernatant was removed. Cell pellet was resuspended gently in 4ml of 4% paraformaldehyde and briefly vortexed at a low speed before being rotated on a rotation spinner for 10 minutes at RT. After fixation, samples were centrifuged and supernatant removed. Cells were resuspended in 2ml of 0.1% BSA in PBS. After resuspension, cells were filtered through a 70μm strainer (Miltenyi Biotec) to filter out any cell clumps. Cells were then centrifuged again, and the supernatant was removed. Cell pellets were then resuspended in 1ml of permeabilization/blocking buffer (0.1% Triton X-100, 1% BSA, 10% normal goat serum (Sigma) in PBS), and incubated on a rotation spinner for 30 minutes at RT. After permeabilization/blocking, cells were centrifuged and the supernatant was removed. Cells were then resuspended in the primary antibodies (1:200) made up in 0.1% BSA in PBS, and incubated on the rotation spinner for 1 hour at RT. After primary antibody incubation, cells were centrifuged, supernatant removed and washed once in 0.1% BSA in PBS. They were then resuspended in the species-specific secondary antibodies (AlexaFluor 488, 647) at a dilution of 1:500 made up in 0.1% BSA in PBS and incubated in the dark on a rotation spinner for 30 minutes. After incubation, cells were centrifuged, supernatant removed and washed once in PBS, followed by incubation with DAPI made up in PBS for 5 minutes. The DAPI + PBS was then removed, followed by one wash in PBS, before being analysed on the flow cytometer.

The samples were run on the LSRii (BD) cell sorter. Scattering was initially used to discard debris as well as cell doublets and larger clumps. The single-cell population was then gated to include DAPI positive only cells (negative control). The gating threshold for measured channels was determined using the control lacking the antibody of interest (Fluorescence minus one (FMO) control), for both channels being recorded. Once the parameters had been set, 10,000 cell events were recorded, and data were processed and analysed on FlowJo (RRID:SCR_008520, https://www.flowjo.com/solutions/flowjo).

### High Performance Liquid Chromatography (HPLC) and sample preparation

The mDA neurons from three independent hiPSC lines were at 3 and 4 weeks of differentiation were incubated in phenol-free “N2B27” medium for 24 hours with and without the presence of 80μM L-Dopa (Sigma D9628). For extracellular metabolites, media after 24 hours was mixed 1:1 in 0.8M ice cold perchloric acid. Samples were incubated on ice for 10 minutes, followed by centrifugation at 12,000g for 5 minutes at 4°C. The supernatant was removed and frozen in dry ice for HPLC analysis. For intracellular metabolites, after 24 hours, the cells were removed and pelleted by centrifugation at 3600 RPM for 5 minutes. The cells were washed once in PBS and lysed on ice using lysis buffer (10mM Tris (pH 7.4), 1mM EDTA, 320mM sucrose in HPLC grade water). Lysate was mixed with 1:1 in 0.8M ice cold perchloric acid and incubated on ice for 10 minutes. The samples were centrifuged at 12,000g for 10 minutes and the supernatant was harvested and frozen for HPLC analysis.

Quantification of neurometabolites (DOPAC, 3-OMD, 5-HIAA, HVA and dopamine) was carried out using reverse phase HPLC and an electrochemical detector following a method by ^59^. The mobile phase (flow rate 1.5ml/min) contained 16% methanol, 20mM sodium acetate trihydrate, 12.5mM citric acid monohydrate, 3.35mM 1-octanesulfonic acid, 0.1mM EDTA disodium and adjusted to pH 3.45 with 12 M hydrochloric acid (HCl). The stationary phase was maintained at 27°C. The detector electrode was set at 450mV and screening electrode at 50mV. 50μl of sample was injected and calculated against a 500nM external standard solution of the 5 compounds of interest made in HPLC grade water acidified with 12M HCl. Peak areas were quantified with EZChrom Elite™ chromatography data system software, version 3.1.7 (https://www.agilent.com/en-us/support/software-informatics/openlab-software-suite/openlab-cds/ezchromelite320) (JASCO UK Ltd., Great Dunmow, UK).

### Electrophysiology

Visualized patch-clamp recordings from cell cultures were performed using an infrared differential interference contrast imaging system and a Multipatch 700B amplifier controlled by pClamp 10.2 software package (Molecular Devices, USA) (RRID:SCR_011323, http://www.moleculardevices.com/products/software/pclamp.html). For the recordings, a neuronal culture on a glass coverslip was placed in a recording chamber mounted on the stage of an Olympus BX51WI upright microscope (Olympus, Japan). The perfusion solution contained the following (in mM): 119 NaCl, 2.5 KCl, 1.3 Na_2_SO_4_, 2.5 CaCl_2_, 26.2 NaHCO_3_, 1 NaH_2_PO_4_, 2 CaCl_2_, 2 MgCl_2_, 22 glucose and was continuously bubbled with 95% O_2_ and 5% CO_2_, pH 7.4. Whole-cell recordings were performed at 32-34°C; the patch-clamp pipette resistance was 3-7 MΩ depending on particular experimental conditions. Series resistance was monitored throughout experiments using a +5 mV step command, cells with very high series resistance (above 25 MΩ) or unstable holding current were rejected. The intracellular pipette solution for voltage-clamp experiments contained (in mM): 120.5 CsCl, 10 KOH-HEPES, 2 EGTA, 8 NaCl, 5 QX-314 Br^-^ salt, 2 Na-ATP, 0.3 Na-GTP. For current-clamp experiments, the intracellular solution contained (in mM): 126 K-gluconate, 4 NaCl, 5 HEPES, 15 glucose, 1 K_2_SO_4_×7 H_2_O, 2 BAPTA, 3 Na-ATP. pH was adjusted to 7.2 and osmolarity adjusted to 295 mOsm. To isolate response of NMDA receptors we added to a perfusion solution: 50 mM picrotoxin, 20 mM NBQX, 1 mM strychnine, 1 mM CGP-55845, 100 mM MCPG, with zero Mg^2+^. To isolate response of GABA_A_ receptors, we added 50 mM APV, 20 mM NBQX, 1 mM strychnine, 1 mM CGP-55845, 100 mM MCPG. All chemicals were purchased from Tocris Bioscience. mDA neurons were tested as a subgroup of the set of generated cultures.

### Live-cell imaging

To measure [Ca^2+^]_c_, cells were loaded with 5μM of Fura-2 AM in HBSS for 30 min at room temperature followed by 2x HBSS washes (Invitrogen). Cells were imaged using epifluorescence on an inverted microscope equipped with a 20x fluorite objective. The cells were excited sequentially at 340 and 380nm using light from a Xenon arc lamp. The emitted fluorescence was measured at 515nm on a cooled camera device (CCD). The fluorescence intensity of the bound and unbound Ca^2+^ was then quantified using ratiometric analysis on Fiji (RRID:SCR_002285, http://fiji.sc).

To measure level of antioxidant, reactive oxygen species (ROS), mitochondrial membrane potential, lysosomal dynamics, calcium uptake, mitochondrial calcium, and cell death, a confocal microscope (ZEISS LSM 710/880 with an integrated META detection system) which has illumination intensity limited to 0.1 −0.2 % of laser output to prevent phototoxicity was used.

The antioxidant level was accessed using glutathione indicator, 50μM Monochlorobimane (mBCI, ThermoFisherScientific) was incubated for 30 minutes and measured at 420 - 550 nm excited by a 405 nm laser. To measure mitochondrial membrane potential, cells were incubated with 25nM tetramethylrhodamine methyl ester (TMRM, ThermoFisherScientific) in HBSS for 40 minutes and then imaging was acquired using Zeiss LSM 880 confocal microscope. The 560 nm laser line was used to excite and it was measured above 560 nm. Approximately 3-5 fields of view with Z projections were taken per sample. To measure lysosomal dynamics, cells were incubated with 50nM LysoTracker™ Deep Red (ThermoFisherScientific), and Hoechst 33342 in HBSS for 40 minutes and then imaged using Zeiss LSM 880 confocal microscope where the 405 nm, and 647 nm laser line were used to excite Hoechst 33342 and LysoTracker^™^ Deep Red, respectively. Approximately 4-5 fields of view with Z projections were taken per sample.

Calcium uptake and dynamics were assessed using the dye Fluo-4 AM (ThermoFisherScientific). Cells were incubated with 5μM Fluo-4 AM in HBSS for 40 min, followed by two HBSS washes. For measuring mitochondrial calcium, cells were incubated with 2μM X-Rhod-1 AM and Fluo-4 AM in HBSS for 40 min, followed by two HBSS washes. For measuring calcium and mitochondrial membrane potential, cells were loaded with Fluo-4 AM and 25nM TMRM for 40 min, followed by two HBSS washes. 25nM of TMRM was then re-added to cells prior to imaging. Live-cell imaging was performed excited by a 480 nm laser and measured at 520 nm. A time-series with 5 second intervals was performed to establish basal fluorescence before 50mM KCl was added to depolarise the membrane and measure fluorescence intensity increase, and recovery.

To measure cell death, live cells were incubated with 100nM SYTOX™ Green Nucleic Acid Stain (ThermoFisherScientific), and the nuclear marker Hoechst 33342 for 40 minutes in HBSS. As SYTOX™ Green is impermeable to live cells, only dead cells were stained, whereas Hoechst 33342 labelled all cells. Cells were imaged using the Zeiss LSM 880 confocal microscope where the 405 nm, and the 488 nm laser line were used to excite Hoechst 33342 and SYTOX Green, respectively. Approximately 4-5 fields of view with Z projections were taken per sample.

To measure ROS, (mainly superoxide) a cooled camera device (CCD) was used and data were obtained on an epifluorescence inverted microscope equipped with a 20x fluorite objective. Cells were loaded with 2μM dihydroethidium (HEt, Molecular Probe) in HBSS. We generated ratios of the oxidised form (ethidium) exited at 530 nm and measured using a 560 nm longpass filter versus the reduced form with excitation at 380 nm measured at 415 - 470 nm.

### Fluorescent false neurotransmitter (FFN) live-cell DAT imaging

To measure the presence and activity of the DAT, we utilized the commercially available fluorescent DAT and VMAT2 substrate FFN102 (Abcam, ab120866). To measure the uptake of the FFN102 dye, a field of view was first found using the brightfield settings on a Zeiss LSM 880 confocal microscope. The cells then had 10μM of the dye in HBSS added and a time-series with an exposure every 5 seconds using the 405 nm laser was started, to measure uptake of the dye into the cells. As a control to confirm specificity, samples in a different well were pre-treated with 5μM of the DAT inhibitor nomifensine (Sigma) for 10 minutes in HBSS. Nomifensine was kept in cell solution also after FFN102 was added. Once the dye had entered cells, the cells were depolarised by the addition of 50mM KCl to observe FFN102 dynamics. Approximately 20 cells were measured per condition/sample and their rate of fluorescence intensity increase was plotted.

### Sample preparation for single molecule localisation microscopy

For single molecule localisation microscopy (SMLM), neurons were grown on glass coated ibidi chambers. Once neurons reached the desired age, they were washed once in PBS, followed by a 15 minute fixation in 4% paraformaldehyde + 0.1% glutaraldehyde (both from Electron Microscopy Services) in PBS at RT. The neurons were then reduced in 0.1% sodium borohydride (Sigma) in PBS for 7 minutes at RT. Cells were then washed 2x in PBS.

To stain cells with aptamer and phalloidin, after fixation and PBS washes, cells were permeabilised with 0.25% triton X-100 in PBS for 10 minutes at RT. The cells were then blocked in blocking solution (0.1% triton X-100, 10% normal goat serum (Abcam), 10% salmon sperm DNA (Thermo Fisher Scientific)) in PBS for 2 hours at RT. The samples were then incubated with 100 nM of the aptamer (sequence: GCCTGTGGTGTTGGGGCGGGTGCGTTATACATCTA) made up in the blocking solution at 4°C overnight. After incubation, cells were washed 1x in PBS and incubated with phalloidin-647 (1:400) (Thermo Fisher Scientific) made up in the blocking solution for 1 hour at RT. Cells were then washed 1x in PBS and either imaged, or incubated with DAPI (1:10000) in PBS for 10 minutes at RT followed by 2x PBS washes before imaging.

### Single molecule localisation microscopy

SMLM was performed on a Nanoimager super-resolution microscope (Oxford Nanoimaging Ltd) equipped with an Olympus 1.4 NA 100x oil immersion super apochromatic objective. To ensure efficient blinking for STORM (AF647-tagged phalloidin), the samples were incubated with a blinking induction buffer (B cubed, ONI). Separately to this, DNA-PAINT was also employed which relies on the addition of an imaging strand (sequence: CCAGATGTAT-CY3B) to the buffer. 1 nM of the imaging strand was added to the B cubed buffer before imaging. The laser illumination angle was set to 51° for all imaging leading to total internal reflection fluorescence (TIRF). AF647-tagged phalloidin was first imaged for 4000-8000 frames using the 640 nm laser (80% power). After this, 4000-5000 frames at 30% power for the 561 nm laser was used to image and super-resolve the aptamer. Both were recorded at a frame-rate of 50 ms. This was done for 2-3 fields of view per line and condition.

### Structured illumination microscopy (SIM)

Samples for SIM were cultured on glass bottom Ibidi 8-well chambers coated with laminin. They were fixed and stained for intracellular markers as described in the ICC section. SIM was performed on an Elyra PS.1 microscope (Zeiss), using a 40x oil objective (EC Plan-Neofluar 40x/1.30 Oil DIC M27). Images were acquired as 15x 0.1 μm Z-planes on a pco.edge sCMOS camera, using 5 grid rotations with the 405 nm (23μm grating period), the 488 nm (28μm grating period) and the 561 nm (34μm grating period) lasers. Images were processed and channels aligned using the automatic settings on the ZEN Black software (Zeiss) (RRID:SCR_018163, http://stmichaelshospitalresearch.ca/wp-content/uploads/2015/09/ZEN-Black-Quick-Guide.pdf).

### Image analysis

ICC images, lysosomal dynamics, and cell death were analysed using Fiji. Live-cell imaging quantification was analysed using the ZEN software by getting the fluorescence intensities of the region of interest for each frame. All graphs and traces were plotted on Prism 8 (GraphPad) (RRID:SCR_002798, http://www.graphpad.com/), with the exception of HEt traces in Figure 6C, which were analysed and plotted using Origin 2018 (Microcal Software Inc., Northampton, MA) (RRID: SCR_014212).

All SMLM analysis was performed using the Oxford Nanoimaging Ltd developed online software, CODI (https://pages.oni.bio/codi-advanced-ev-characterization-made-simple). The super-resolved images are displayed as the output of all super-resolved single-molecule localisations from the entire frame acquisition. Once super-resolution images were uploaded to the software, initially an inbuilt drift-correction was performed in order to correct single-molecule localisations in case the sample drifted in the x or y direction during acquisition. After the drift correction, the number of frames was changed to only include the frames where the relevant fluorophore was imaged. Each localisation was fitted to a 2D Gaussian distribution and any of those with a standard deviation larger than 250 nm were removed. Finally, any localisations with a precision lower than 20 nm were discarded. Once the filtered super-resolved image was generated, density-based spatial clustering of applications with noise (DBSCAN) was performed on the resulting images. Each cluster in DBSCAN needed to have at least 15 localisations, and each localisation had to be within 60 nm of each other. This was to remove any non-specific aptamer binding, and to only detect aggregates that were quite spatially confined.

### Statistical analysis

Statistical analysis was performed on Prism 8. To compare two individual groups, a unpaired, two-tailed t-test was used to generate a p value. When comparing more than two individual groups, an ordinary one-way ANOVA was used with a post hoc Tukey test for multiple comparisons between groups. When comparing two individual variables, an ordinary two-way ANOVA was performed, with a correction of the False Discovery Rate or the Tukey’s range test for multiple comparisons between groups. A p value below 0.05 was considered to be statistically significant. Results are represented as means ± standard error of the mean (SEM), or standard deviation (SD) where stated in the figure legends. The number of hiPSC lines, number of cells, and the number of neuronal inductions used for each experiment is stated in the respective figure legend. The sizes of the sample for each experiment was selected to ensure that the technical (number of cells, numbers of fields of view, and number of coverslips), and biological (number of hiPSC lines, and number of neuronal inductions) variation was adequately captured, and is listed in the respective figure legends.

