## Supplementary material for "Protein aggregation and calcium dysregulation are the earliest hallmarks of synucleinopathy in human midbrain dopaminergic neurons"

**Supplementary Table 1. hiPSC lines used in this study**

| hiPSC line | Mutation | Age of donor | Sex of donor | Source | RRID |
|---|---|---|---|---|---|
| No disease Ctrl 1 | None | 78 | Male | Reprogrammed in house EDi046-A | |
| No disease Ctrl 2 | None | 51 | Male | Cedars Sinai iPSC Core Repository CS0002iCTR-nxx | |
| No disease Ctrl 3 | None | Unknown | Female | Thermo Fisher Scientific A18945 | CVCL_RM92 |
| No disease Ctrl 4 | None | 64 | Male | Coriell ND41866 | CVCL_Y838 |
| A53T 1 & isogenic control | *SNCA* A53T | 54 | Female | StemBANCC SFC828 (STBCi019-A) | CVCL_RB71 |
| A53T 2 | *SNCA* A53T | 57 | Male | StemBANCC SFC829 (STBCi023-C) | CVCL_RB78 |
| *SNCA* x3 | *SNCA* locus x3 | Unknown | Female | StemBANCC SFC831 (STBCi024-C) | CVCL_RB81 |
| *SNCA* x3 & isogenic control | *SNCA* locus x3 | Unknown | Female | As reported [51] EDi001-B | CVCL_ZA47 |

**Supplementary Table 2. List of antibodies used in this study.**

| Protein | Company | Catalogue | RRID | Species | Dilution |
|---|---|---|---|---|---|
| LMX1A | Abcam | ab139726 | AB_2827684 | Rabbit | 1:500 |
| FOXA2 | Santa-Cruz Biotechnology | sc-374376 | AB_10989742 | Mouse | 1:100 |
| OTX2 | R & D Systems | AF1979 | AB_2157172 | Goat | 1:500 |
| TH | Abcam | ab137869 | AB_2801410 | Rabbit | 1:500 (ICC) 1:200 (Flow) |
| TH | Abcam | ab76442 | AB_1524535 | Chicken | 1:500 |
| TUJ1 | Biolegend | 801201 | AB_2313773 | Mouse | 1:1000 (ICC) |
| Beta III Tubulin | Abcam | ab41489 | AB_727049 | Chicken | 1:500 (ICC) 1:200 (Flow) |
| MAP2 | Abcam | ab11267 | AB_297885 | Mouse | 1:500 |
| GIRK2 | Alomone Labs | APC-006 | AB_2040115 | Rabbit | 1:400 |
| Total alpha-synuclein | Abcam | ab138501 | AB_2537217 | Rabbit | 1:200 |
| Filament alpha-synuclein | Abcam | ab209538 | AB_2714215 | Rabbit | 1:50-1:100 |
| Tomm20 | Santa-Cruz | sc-17764 | AB_628381 | Mouse | 1:1000 |
| LC3B | Cell Signaling Technology | #3868 | AB_2137707 | Rabbit | 1:300 |
| LC3B | Abcam | ab48394 | AB_881433 | Rabbit | 1:2000 |
| Phosphorylated alpha-synuclein | Abcam | ab51253 | AB_869973 | Rabbit | 1:1000 |
| Goat pAb anti-Mouse IgG Alexa Fluor 488 | Abcam | ab150113 | AB_2576208 | Goat | 1:500 |
| Goat pAb anti-Mouse IgG Alexa Fluor 555 | Abcam | ab150114 | AB_2687594 | Goat | 1:500 |
| Goat pAb anti-Mouse IgG Alexa Fluor 647 | Abcam | ab150115 | AB_2687948 | Goat | 1:500 |
| Goat pAb anti-Rabbit IgG Alexa Fluor 488 | Abcam | ab150077 | AB_2630356 | Goat | 1:500 |

| | | | | | |
|---|---|---|---|---|---|
| Goat pAb anti-Rabbit IgG Alexa Fluor 555 | Abcam | ab150078 | AB_2722519 | Goat | 1:500 |
| Goat pAb anti-Rabbit IgG Alexa Fluor 647 | Abcam | ab150079 | AB_2722623 | Goat | 1:500 |
| Goat pAb anti-Chicken IgG Alexa Fluor 647 | Abcam | ab150171 | AB_2921318 | Goat | 1:500 |

**Supplementary Table 3. List of TaqMan$^{TM}$ Gene Expression probes used in this study.**

| Gene | Catalogue Number |
|---|---|
| FOXA2 | Hs00232764_m1 |
| LMX1A | Hs00898455_m1 |
| EN1 | Hs00154977_m1 |
| TH | Hs00165941_m1 |
| Nurr1 | Hs00428691_m1 |
| DAT | Hs00997374_m1 |
| GIRK2 | Hs01040524_m1 |
| SNCA | Hs00240906_m1 |
| GAPDH | Hs03929097_g1 |

**Supplementary table 5: Protocols used in this study deposited on protocols.io**

| Protocol | DOI |
|---|---|
| Human iPSC cell culture | dx.doi.org/10.17504/protocols.io.81wgby7dnvpk/v1 |
| Generation of midbrain dopaminergic neurons | dx.doi.org/10.17504/protocols.io.x54v9j7ezg3e/v1 |
| Single-cell RNA-seq | dx.doi.org/10.17504/protocols.io.6qpvr4dxpgmk/v1 |
| Live-cell imaging: Calcium | dx.doi.org/10.17504/protocols.io.3byl4j7e8lo5/v1 |
| Live-cell imaging: Mitochondria membrane potential | dx.doi.org/10.17504/protocols.io.e6nvwj997lmk/v1 |
| Live-cell imaging: Cell death | dx.doi.org/10.17504/protocols.io.n2bvj8yywgk5/v1 |
| Live-cell imaging: Reactive oxygen species (Superoxide) | dx.doi.org/10.17504/protocols.io.5qpvor55zv4o/v1 |
| Fluorescent false neurotransmitter (FFN) DAT imaging | dx.doi.org/10.17504/protocols.io.36wgqj55xvk5/v1 |
| Quantitative polymerase chain reaction (qPCR) | dx.doi.org/10.17504/protocols.io.36wgqj5rkvk5/v1 |
| Immunocytochemistry | dx.doi.org/10.17504/protocols.io.q26g74w79gwz/v1 |

| | |
|---|---|
| Sample preparation for single molecule localisation microscopy and iSIM and microscopy | dx.doi.org/10.17504/protocols.io.3byl4j77olo5/v1 |
| Flow Cytometry | dx.doi.org/10.17504/protocols.io.kqdg3956eg25/v1 |
| High Performance Liquid Chromatography (HPLC) and sample preparation | dx.doi.org/10.17504/protocols.io.yxmvm2ko5g3p/v1 |
| Electrophysiology | dx.doi.org/10.17504/protocols.io.4r3l274mjg1y/v1 |

Supplementary Figure 1

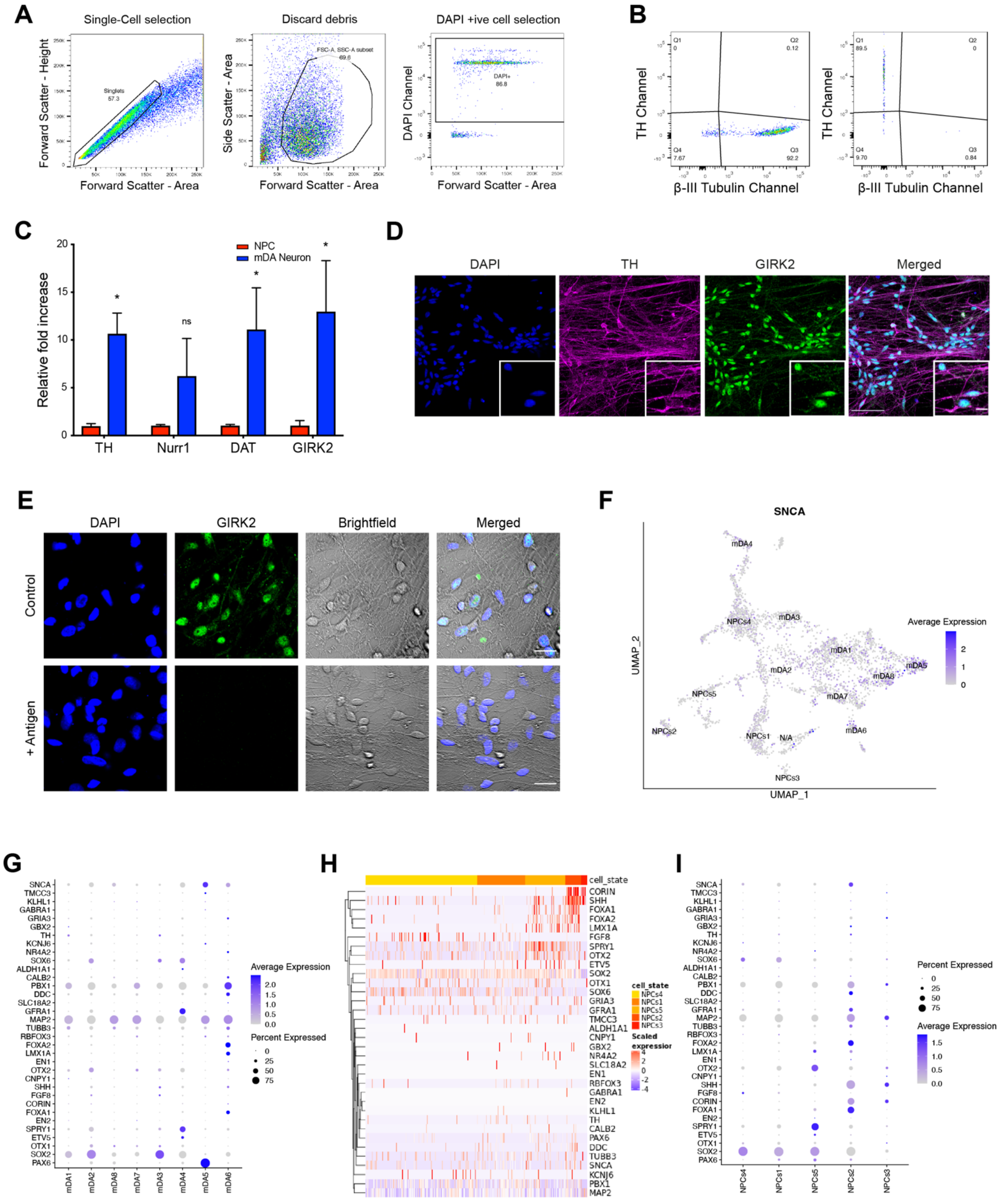

Supplementary Figure 2

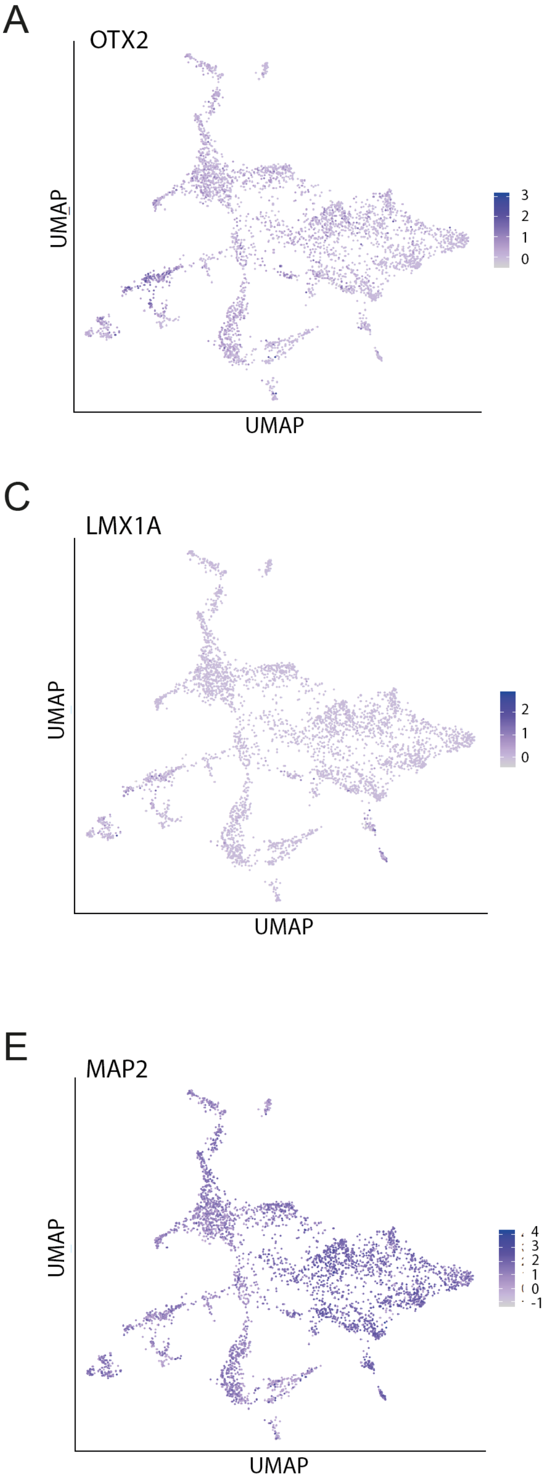

Supplementary Figure 3

A

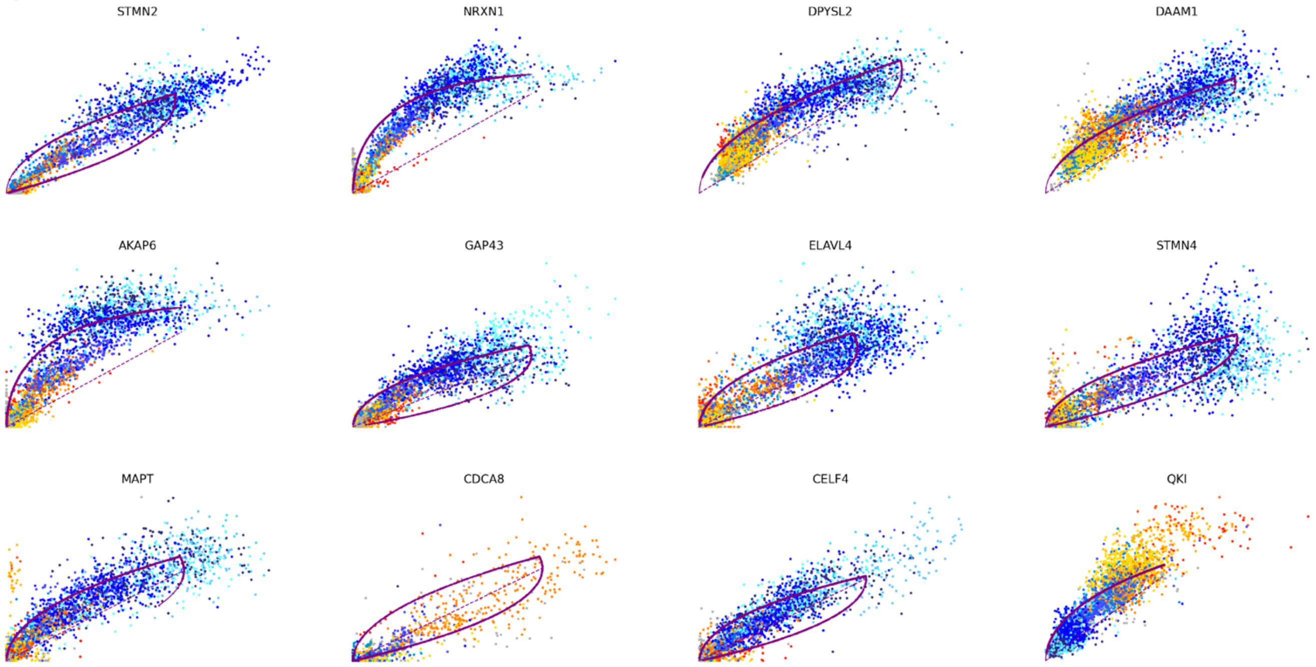


B

| | N/A | NPCs1 | NPCs2 | NPCs3 | NPCs4 | NPCs5 | mDA1 | mDA2 | mDA3 | mDA4 | mDA5 | mDA6 | mDA7 | mDA8 |
|---|---|---|---|---|---|---|---|---|---|---|---|---|---|---|
| **0** | CDCA8 | DAAM1 | DPYSL2 | NDC80 | QKI | DAAM1 | NRXN1 | DAAM1 | DAAM1 | DAAM1 | STMN2 | STMN2 | STMN2 | STMN2 |
| **1** | TM4SF1 | DPYSL2 | DAAM1 | PBK | MEST | MAP1B | AKAP6 | NRXN1 | DCX | DPYSL2 | DPYSL2 | AKAP6 | NRXN1 | NRXN1 |
| **2** | TOP2A | FNBP1L | GAP43 | SFRP1 | VCAN | MEST | DPYSL2 | MAPT | CSRNP3 | TM4SF1 | DAAM1 | GAP43 | DPYSL2 | GAP43 |
| **3** | CAVIN1 | SYT1 | ELAVL4 | VCAN | HMGA2 | CADM2 | GAP43 | STMN2 | FNBP1L | FNBP1L | CSRNP3 | DAAM1 | AKAP6 | DAAM1 |
| **4** | AURKB | QKI | NCAM1 | TPX2 | MAP1B | PON2 | MAPT | DPYSL2 | KIF5C | CAV1 | GAP43 | MEST | CSRNP3 | AKAP6 |

C

| | N/A | NPCs1 | NPCs2 | NPCs3 | NPCs4 | NPCs5 | mDA1 | mDA2 | mDA3 | mDA4 | mDA5 | mDA6 | mDA7 | mDA8 |
|---|---|---|---|---|---|---|---|---|---|---|---|---|---|---|
| **N/A** | 0 | 0.28 | 0 | 0.017 | 0 | 0 | 0 | 0 | 0 | 0 | 0 | 0 | 0 | 0 |
| **NPCs1** | 0 | 0 | 0 | 0 | 0 | 0 | 0 | 0.42 | 0 | 0 | 0 | 0 | 0 | 0 |
| **NPCs2** | 0 | 0 | 0 | 0 | 0 | 0 | 0 | 0.29 | 0 | 0 | 0 | 0 | 0 | 0 |
| **NPCs3** | 0 | 0 | 0 | 0 | 0 | 0 | 0 | 0 | 0 | 0 | 0.18 | 0.15 | 0 | 0 |
| **NPCs4** | 0 | 0 | 0 | 0 | 0 | 0 | 0 | 0.47 | 0 | 0 | 0 | 0 | 0 | 0 |
| **NPCs5** | 0 | 0 | 0 | 0 | 0 | 0 | 0 | 0.33 | 0 | 0 | 0 | 0 | 0 | 0 |
| **mDA1** | 0 | 0 | 0 | 0 | 0 | 0 | 0 | 0 | 0 | 0 | 0 | 0 | 0.32 | 0.35 |
| **mDA2** | 0 | 0 | 0 | 0 | 0 | 0 | 0 | 0 | 0 | 0 | 0 | 0 | 0 | 0 |
| **mDA3** | 0 | 0 | 0 | 0 | 0 | 0 | 0.32 | 0.3 | 0 | 0 | 0 | 0 | 0 | 0 |
| **mDA4** | 0 | 0 | 0 | 0 | 0 | 0 | 0 | 0.39 | 0 | 0 | 0 | 0 | 0 | 0 |
| **mDA5** | 0 | 0 | 0 | 0 | 0 | 0 | 0 | 0 | 0 | 0 | 0 | 0 | 0 | 0 |
| **mDA6** | 0 | 0 | 0 | 0 | 0 | 0 | 0 | 0 | 0 | 0 | 0 | 0 | 0 | 0 |
| **mDA7** | 0 | 0 | 0 | 0 | 0 | 0 | 0 | 0 | 0 | 0 | 0 | 0 | 0 | 0 |
| **mDA8** | 0 | 0 | 0 | 0 | 0 | 0 | 0 | 0 | 0 | 0 | 0 | 0 | 0 | 0 |

Supplementary Figure 4

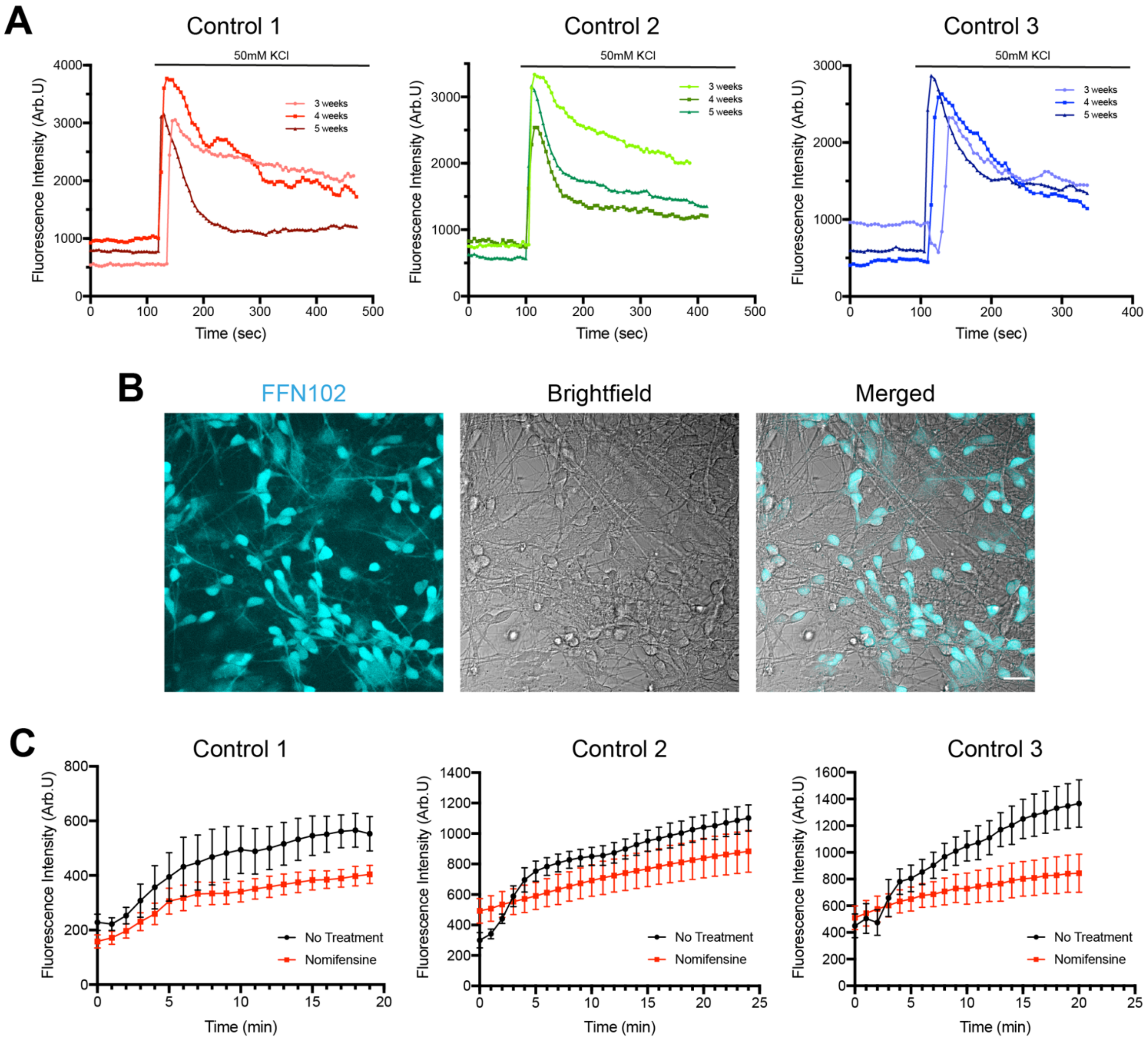

Supplementary Figure 5

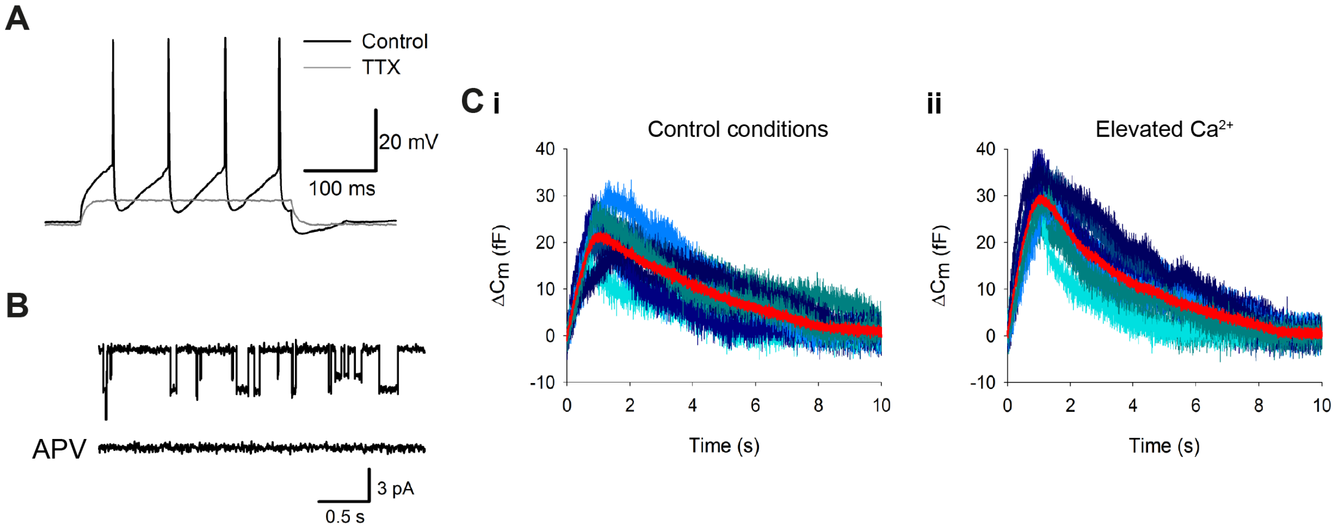

Supplementary Figure 6

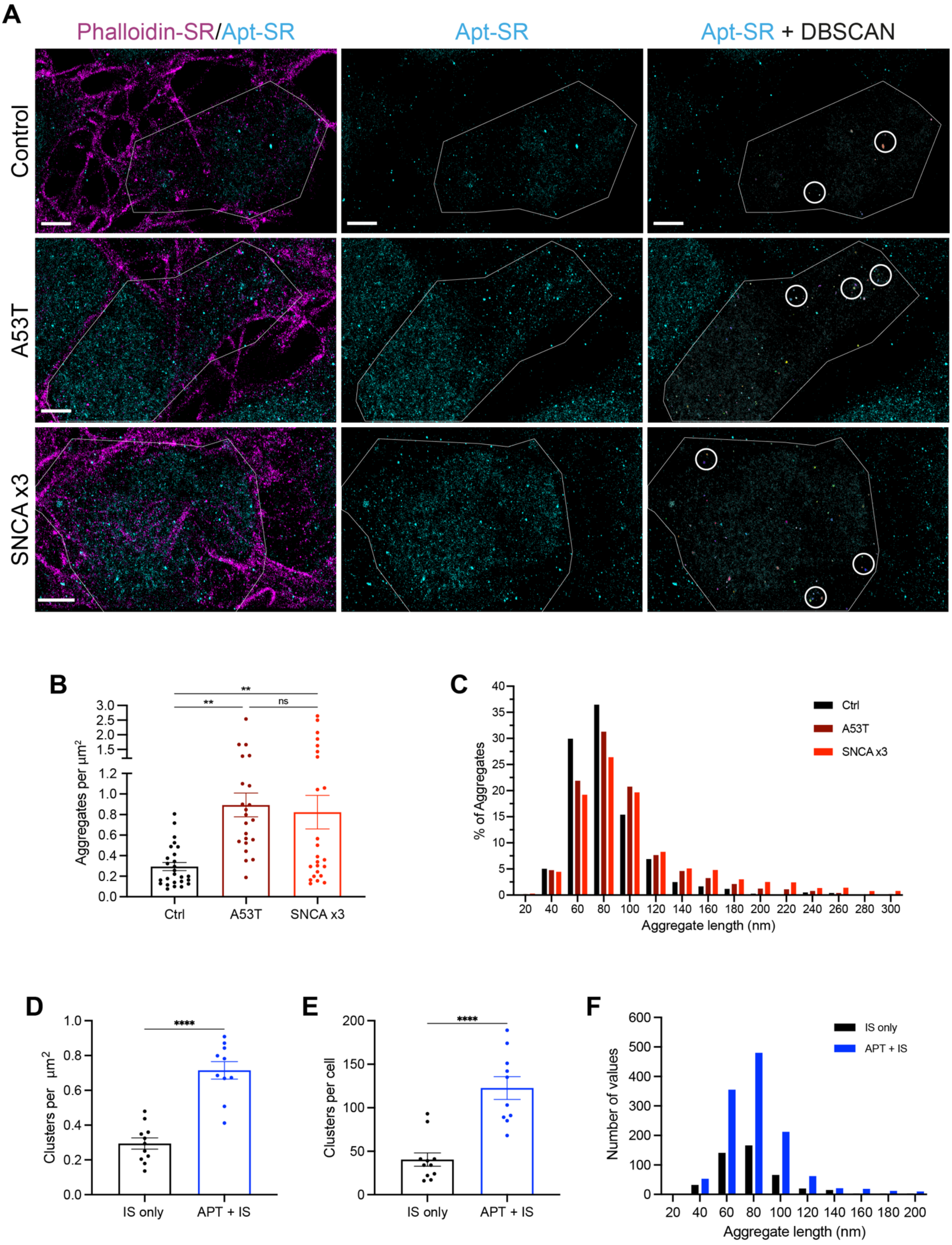

Supplementary Figure 7

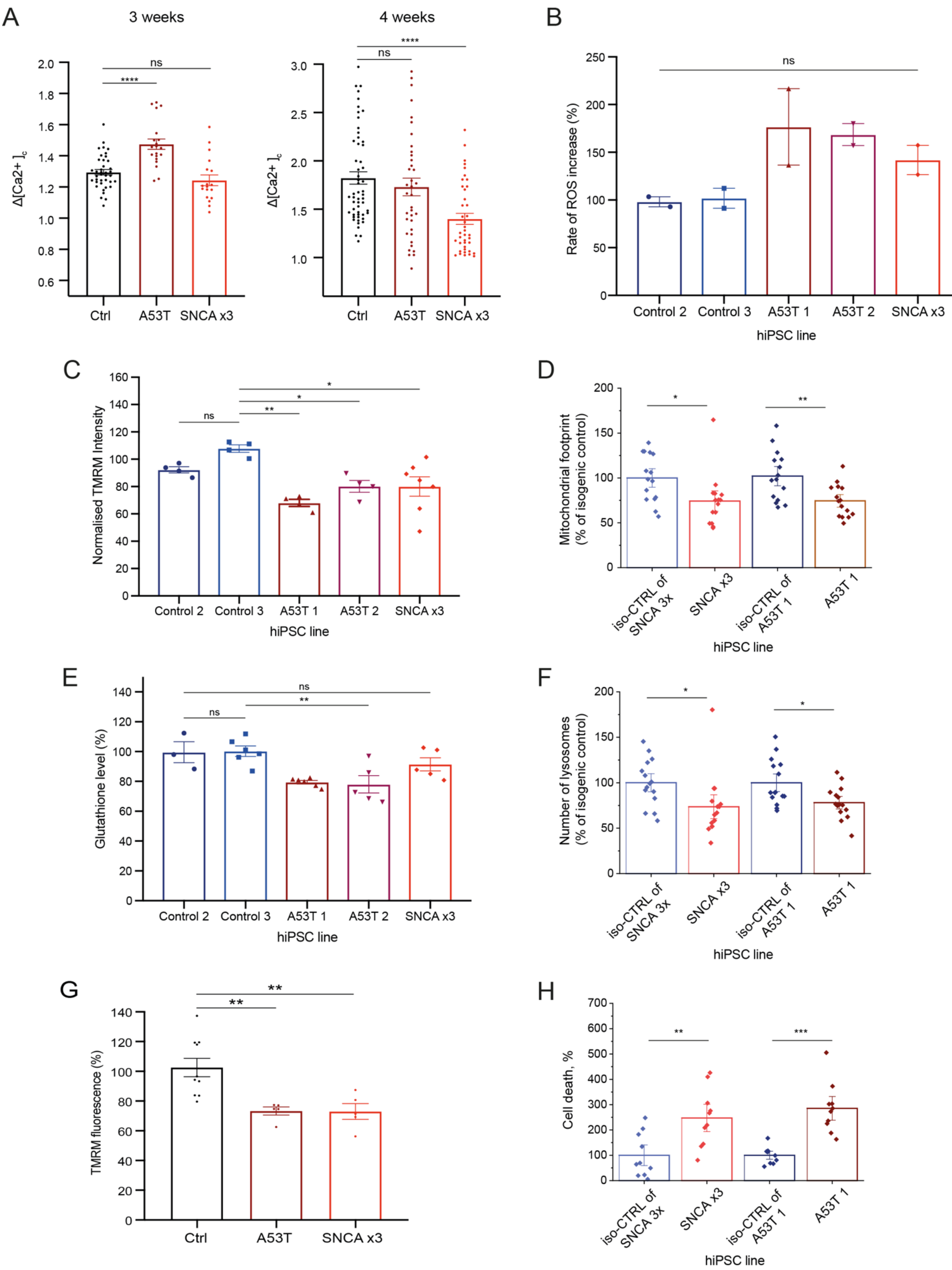

### Supplementary Figure legends

#### SFigure 1 – Characterisation of mDA neurons.

**(A)** Dot plots showing the selection criteria for flow cytometry analysis. Using the forward scatter, only single cells were selected. Next, in the middle dot plot, cluster was selected discarding any debris. Finally, only the DAPI positive cells were selected. **(B)** Dot plots showing the thresholding gating for the channels measured. Thresholding gates were determined using the fluorescence minus one (FMO) control. In the first dot plot, the TH gate threshold was set by measuring the intensity in the sample with all immunolabelling except TH. The same was done for the β-III Tubulin gate threshold in the right dot plot **(C)** Quantitative PCR showing an up-regulation of mRNA of mature mDA markers *TH, Nurr1*, *DAT*, and *GIRK2* relative to mDA NPCs (n = 3 different lines, 3 neuronal inductions, ns $p > 0.05$, * $p < 0.05$, two-way ANOVA). Values plotted as ±SEM. **(D)** Representative ICC images showing expression of TH and GIRK2. Scale bar = 50μm. Smaller image depicts a zoomed-in version of the image showing co-expression of TH and GIRK2. Scale bar = 10μm. **(E)** GIRK2 Antibody specificity was confirmed using the control antigen. Both nuclear and cytosolic signal using a GIRK2 antibody was abolished on applying the antibody incubated with the control GIRK2 antigen. The top panel shows GIRK2 using the antibody as normal. The lower panel shows GIRK2 immunolabelling is abolished when the antibody was incubated with the antigen (GIRK2 sequence). Scale bar = 20μm. **(F)** A feature plot showing the expression of *SNCA* in all the clusters identified through single-cell RNA-seq at 4 weeks of differentiation. **(G)** A dot-plot showing the gene expression profile of the mDA clusters (mDA1-8) identified in single-cell RNA-seq. Genes corresponding to mDA NPC markers and mDA neuron markers are shown. Expression of the genes is coloured based on the expression fold difference and plotted based on the percentage of cells in that cluster expressing the gene. **(H)** A heat map showing the expression of genes in the clusters identified as mDA NPCs (NPCs1-5). Each line represents a cell from that cluster. **(I)** A dot-plot showing the gene expression profile of the mDA NPC clusters. Genes corresponding to mDA NPC markers and mDA neuron markers are shown.

#### SFigure 2 – RNAseq UMAP plots for the key markers of mDA neurons.

UMAP feature plots showing percentage gene expression for the key markers (**A)** OTX2, **B:** FOXA2, **C:** LMX1A, **D:** LMX1B, **E:** MAP2 and **F:** TH) of mDA neurons identified from single-cell RNA-seq after 4 weeks of differentiation.

**SFigure 3 – RNA-velocity on mDA neurons.**

**(A)** Phase portraits of some of the driver genes identified in each cluster. Blue dots represent mDA neuron clusters, yellow dots represent NPC clusters. Driver genes for neuronal clusters have higher amounts of blue cells at the top of the plot (*STMN2*) whereas driver genes for NPC clusters (*QKI*) have a higher amount of cells at the top. Lines represent the "steady-state" ratio, i.e. the ratio of unspliced to spliced mRNA abundance which is in a constant transcriptional state. *QKI* for example is expressed (steady-state expression) more in NPCs, while it is not expressed in mDA neurons. *ELAVL4* on the other hand is expressed at a steady-state in mDA neurons and is up-regulated in NPCs. **(B)** A table showing the top 5 driver genes for each cluster identified through RNA-velocity. **(C)** PAGA analysis visualised in Figure 2C in a table showing the numeric expression of the transition confidences between clusters.

**SFigure 4 – $Ca^{2+}$ and DAT functionality in mDA neurons.**

**(A)** Traces showing the Fluo-4 mean fluorescence intensity for each hiPSC control line before and after the addition of 50mM KCl at different weeks of differentiation (n = 15 cells per trace). Values plotted as the mean of all cells per time point of differentiation**. (B)** Live-cell imaging picture of cells after a 30-minute incubation with FFN, showing the dye enters most mDA neurons. Scale bar = 20µm. **(C)** Traces showing the fluorescence intensity of FFN inside cells to measure the uptake of the dye in each hiPSC line tested, in the absence, or presence of DAT inhibitor nomifensine (n = 15-20 cells per condition). Values plotted as ±SD.

**SFigure 5 – SNCA PD mDA neuron functionality and pathology.**

**(A)** APs triggered by step current injection in SNCA x3 mDA neurons at day 30 of differentiation. 1 mM tetrodotoxin (TTX) fully suppresses APs, confirming involvement of voltage-gated sodium channels. **(B)** Single-channel openings of NMDA receptors in an outside-out patch excised from SNCA x3 mDA neurons at day 30 of differentiation. Top trace: application of 10 mM glutamate + 10 mM glycine triggers single-channel openings with two different conductance states. Bottom trace: 50 mM APV fully suppresses the effect of glutamate + glycine. **(C)** Changes in whole-cell membrane capacitance evoked by stimulation series confirm elevated intensity of vesicle release in SNCA x3 mDA neurons at day 70 of differentiation. Shadows of blue, high noise: 10 consequent individual traces; superficial red trace with low noise: averaged trace. **i)** control. **ii)** Elevated $Ca^{2+}$ magnifies the effect of external stimulation on membrane capacitance; confirming involvement of presynaptic $Ca^{2+}$-dependent mechanism of vesicle release.

**SFigure 6 – Oligomer detection in mDA neurons using SMLM.**

**(A)** SMLM images showing phalloidin in magenta and the aptamer in cyan in 1w mDA neurons. White line shows the ROI of the cell identified. Middle panel shows only the SMLM aptamer. Final panel shows oligomers identified by DBSCAN (in different colours) with example clusters highlighted in circles. Scale bar = 2µm. **(B)** Graph showing the number of aggregates normalised to the area in each cell expressed as aggregate per $µm^2$. **(C)** Each aggregate plotted as a percentage of the total population (y-axis) showing the length of each aggregate binned in 20 nm intervals. **(D)** Graph showing the aggregates per $µm^2$ of each cell when samples were imaged with only the imaging strand (IS) or with prior incubation with the aptamer (n = 1 control lines, 2 fields of view per condition 4-7 cells per field of view, **** p < 0.0001 unpaired t-test). **(E)** Graphs showing the clusters per cell when imaged with only the IS or with prior incubation with the aptamer (**** p < 0.0001 unpaired t-test). **(F)** Histogram showing the length of each cluster binned in 20 nm intervals in IS only and the with incubation of the aptamer (n = 400-1300 aggregates per condition). All values plotted as ±SEM.

**SFigure 7 – SNCA PD mDA neuron functionality and pathology.**

**(A)** Quantification of the change in cytosolic $Ca^{2+}$ amplitude ($\Delta[Ca^{2+}]_c$) after KCl addition in 3 and 4 week old neurons (n = 15-20 cells per line, 2 control hiPSC lines, 1 A53T line, 1 SNCA x3 line, 1 neuronal induction, ns p > 0.05, **** p < 0.0001, one-way ANOVA). **(B)** Relative increase in ROS based on HEt ratiometric fluorescence plotted out for each line tested (n = 2 coverslips per line, ns p > 0.05, one-way ANOVA). **(C)** The normalised TMRM fluorescence intensity plotted out for each line tested using healthy control (n = 3-5 fields of view per line, across 2 coverslips per line, ns p > 0.05, ** p < 0.005, one-way ANOVA). **(D)** The normalised mitochondrial fingerprint measurement  plotted out for each line tested using the isogenic control lines for each mutant (n = 10-15 fields of view per line, 1 SNCA x3 and the isogenic control lines. 1 A53T and the isogenic control lines, * p < 0.05, ** p < 0.005, one-way ANOVA). **(E)** Relative percentage of endogenous glutathione levels based on MCB fluorescence plotted out for each line tested (n = 3-6 fields of view per line; 2 coverslips per line, ns p > 0.05, ** p < 0.005, one-way ANOVA). **(F)** The normalised number of LysoTracker™ spot  plotted out for each line to  tested usingthe isotnice control lines for each mutant (n = 10-15 fields of view per line, 1 SNCA x3 and the isogenic control lines. 1 A53T and the isogenic control lines * p < 0.05, one-way ANOVA). **(G)** Quantification of normalised TMRM fluorescence intensity after the addition of KCl in 6 week old mDA neurons (n = 5-10 cells per line, 1 control line, 1 A53T line, 1 SNCA x3 line, 1 neuronal induction, ** p < 0.005, one-way ANOVA). **(H)** Relative percentage of cell death for each line tested uinsg the isogenic control lines (n = 8-10 fields of view per line, 1 SNCA x3 and the isogenic control lines. 1 A53T and the isogenic control lines, ** $p < 0.005$, *** $P < 0.0005$ one-way ANOVA). All values plotted as ±SEM.
